# The Ithildin library for efficient numerical solution of anisotropic reaction-diffusion problems in excitable media

**DOI:** 10.1101/2024.05.01.592026

**Authors:** Desmond Kabus, Marie Cloet, Christian Zemlin, Olivier Bernus, Hans Dierckx

## Abstract

Ithildin is an open-source library and framework for efficient parallelized simulations of excitable media, written in the C++ programming language. It uses parallelization on multiple CPU processors via the message passing interface (MPI). We demonstrate the library’s versatility through a series of simulations in the context of the mono-domain description of cardiac electrophysiology, including the S1S2 protocol, spiral break-up, and spiral waves in ventricular geometry. Our work demonstrates the power of Ithildin as a tool for studying complex wave patterns in cardiac tissue and its potential to inform future experimental and theoretical studies. We publish our full code with this paper in the name of open science.

**Author summary:** We present Ithildin, an open-source library for reaction-diffusion systems such as the electrical waves in cardiac tissue controlling the heart beat. We demonstrate the versatility of Ithildin by example simulations in various tissue models and geometries, from simple 2D simulations to detailed ones in ventricular geometry. Our simulations highlight key features of Ithildin, such as recording pseudo-electrograms or filament trajectories. We hope that our work will contribute to the growing understanding of cardiac electrophysiology and inform future experimental and theoretical studies.

## 1 Introduction

In the *Lord of the Rings*, Ithildin is an Elven substance that reveals a hidden gateway to an otherwise inaccessible world after a spell is cast [1]. With the Ithildin framework, named after this substance, we want to open up new gateways in the numerical simulation of reaction-diffusion systems, such as the electrical activation patterns in the heart.

Our motivation to write a reaction-diffusion solver comes from the numerical study of electrical patterns inside the heart [2]. These patterns, which are incompletely understood, are a main cause of death and even as a chronic disease, they complicate people’s lives. In the past decades, computer models of arrhythmia have allowed mechanistic insight in the origin and control of arrhythmias [3]. On the longer term, it is thought that digitized versions of patients’ hearts could help offer better diagnostics and planning of procedures [4]. In view of open science, we have decided to share the code that has been steadily developed in our group since 2007 with the scientific community.

A flowchart outlining the functionality of Ithildin can be found in fig. 1. Ithildin is designed to comply with the 2011 version of the ISO-C++ standard [5], but it compiles with all newer versions, including the current 2023 ISO-C++ standard [6–9]. The software facilitates forward Euler and Runge-Kutta finite-difference solutions for reaction-diffusion systems in *N* -dimensional space, such as the mono-domain equation for cardiac electrophysiology, with specified boundary conditions [2].

**Figure 1.**
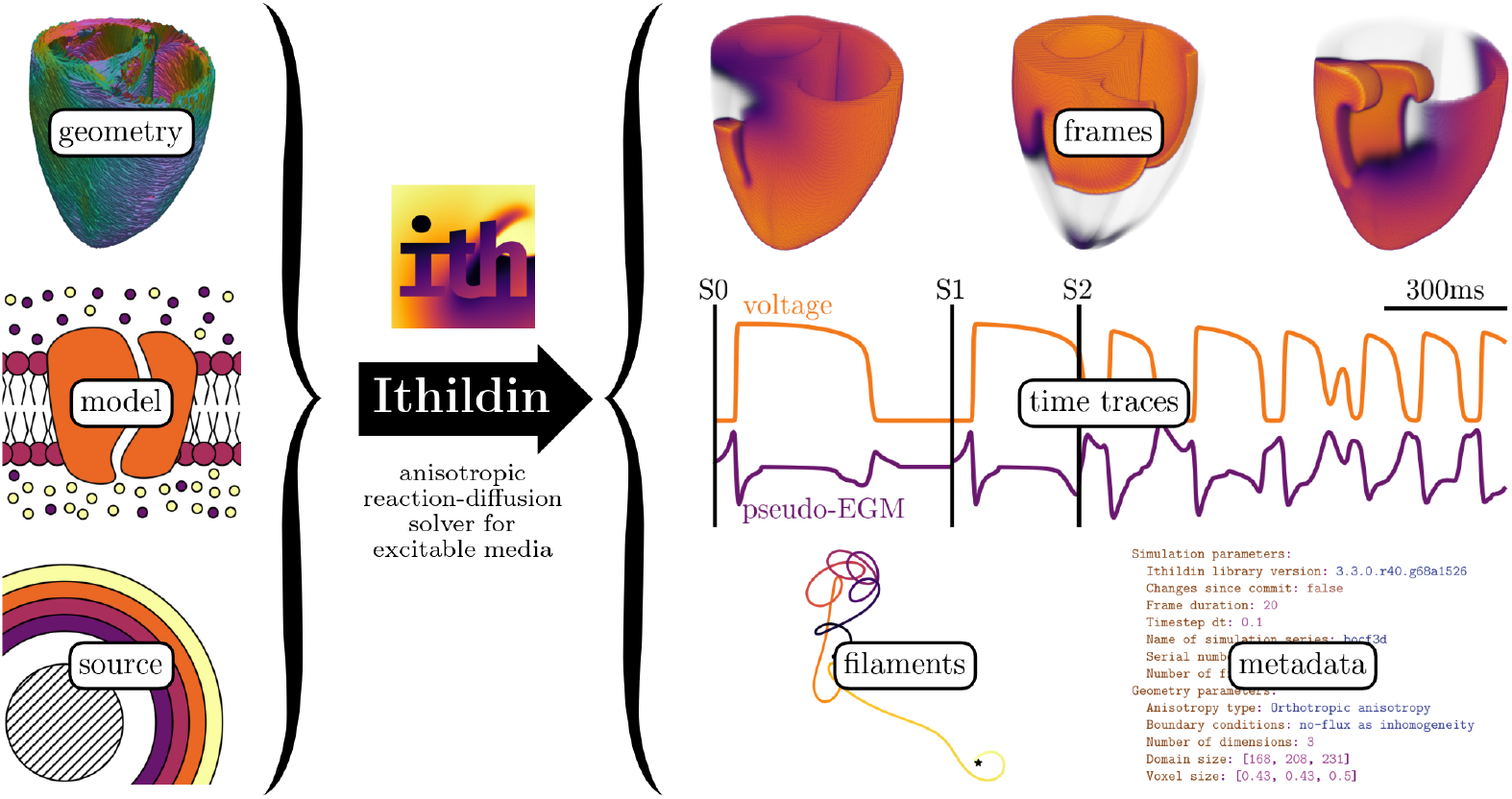
Ithildin can be used to solve reaction-diffusion problems in excitable media. Required inputs for the software are: the diffusion tensor and geometry of a medium—such as the heart muscle, a reaction term—the so-called model, and source terms—typically a stimulation protocol. Ithildin can then calculate the evolution of the model variables in the medium over time. During calculation, Ithildin records relevant spatio-temporal data and metadata, as well as detecting rotor cores as filaments. The data visualized here are taken from several simulations which will be discussed below.

The framework offers quick computation through CPU parallelization using OpenMPI [10]. It also boasts decent documentation of available features, made accessible through Doxygen [11]. Ithildin writes easy-to-parse YAML log files to document simulation setups [12]. Additionally, it allows convenient output of frames of recorded variables at regular intervals in the form of NumPy NPY files [13]. The software supports easy and powerful post-processing with the Python module for Ithildin [14], including integration with Scientific Python [15], 2D visualization with Matplotlib [16], and 3D visualization with ParaView [17].

Ithildin also allows the recording of pseudo-electrograms (EGMs) and state variables at full numerical time resolution, as well as the tracking of filaments, which represent the instantaneous rotation axes of rotors. The software features a flexible setup for in-silico experiments, also called simulations, through a simple class-based C++ interface.

Various types of geometries are implemented, ranging from a simple 1D cable and spirals in 2D tissue to whole-heart geometry and even 4D hyperspace. The space can be partitioned to use multiple cell models in the same experiment via `Model_multi`. Realistic stimulation protocols can be added as `Stimulus` objects and may be started by a `Trigger`. Ithildin also includes a logging system with minimal impact on computation speed and various levels of verbosity.

In this paper, we provide an overview of this framework, guiding the reader through its components. Results from several in-silico experiments are presented as the main components of Ithildin are introduced. The details of these so-called simulations are outlined towards the end of this paper in sec. 5, along with a tabular overview in tbl. 4.

## 2 Essential numerical methods

### 2.1 Reaction-diffusion system

The diffusion of electrical signals in the cardiac tissue is modeled using the diffusion tensor, denoted as ***D***, in the reaction-diffusion equations. This tensor encapsulates the spatial orientation of fibers in the medium and the effects of inhomogeneities on signal propagation. The core equation governing the evolution of the state variable vector *<u>u</u>* is the reaction-diffusion equation:

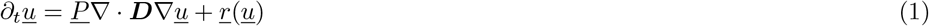

Here, *<u>u</u>* is the state variable vector and *<u>r</u>*(*<u>u</u>*) accounts for the reaction term and is called the *model*. We refer to the first component of *<u>u</u>* as *u*, which for electrophysiological models is the transmembrane voltage *V*_m_ or a rescaled version of it, see also sec. 4.2. For two-variable models, the second component of *<u>u</u>* is often referred to as the restitution variable *v*. In the term representing diffusion, ***D*** is determined by the geometry of the medium and the presence of inhomogeneities, see sec. 4.1. Different notation is used to distinguish between vectors ***x*** and matrices ***D*** in physical space in bold font, and underlined vectors *<u>u</u>* and matrices *<u>P</u>* with respect to state variables. We use lowercase letters for vectors and uppercase for matrices. An overview of the most relevant quantities in Ithildin is given in tbl. 1.

**Table 1.**
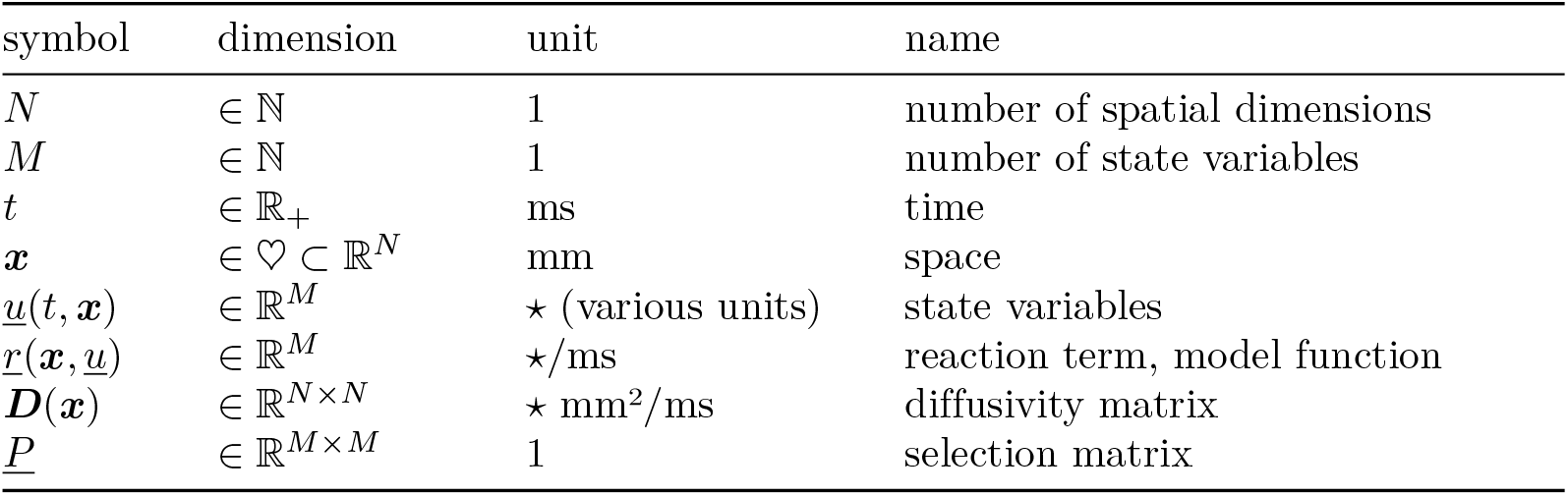
Quantities in the reaction-diffusion problem

Ithildin obtains approximate solutions of the reaction-diffusion equation (eq. 1) via a finite-differences approach: Time *t* and space ***x*** are discretized on a grid and values *<u>u</u>*(*t*, ***x***) are associated with the vertices of this grid. We choose a fixed temporal resolution, the time step Δ*t*, and constant spatial grid spacing Δ***x***. The values *<u>u</u>*(*t* + Δ*t*, ***x***) at a subsequent time-step are computed based on the previous ones, according to discretized versions of the governing equations, i.e., the reaction-diffusion equation (eq. 1), together with boundary and initial conditions.

### 2.2 Time integration

Starting from an initial state, the state variable vector *<u>u</u>* is integrated over time using a so-called time stepping scheme leading to an approximate solution of the reaction-diffusion system using finite differences. Ithildin implements two main stepping schemes to choose from: forward Euler and the classic Runge-Kutta method (RK4) [18, 19].

Defining *<u>f</u>* as the right hand side of the reaction-diffusion equation (eq. 1), the forward Euler method takes the form [19]:

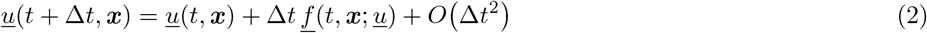

This method is the default time integration scheme in Ithildin. Despite its numerical error being of order *O* (Δ*t*^2^), with a sufficiently small time step, the accuracy of the Euler method is adequate for our use cases.

Due to the Courant-Friedrichs-Lewy condition (CFL), a stability criterion for the integration of the reaction-diffusion equation, Δ*t* needs to be chosen sufficiently small [20]. Ithildin automatically chooses an appropriate time step based on the CFL condition for the different supported geometries, cf. sec. 4.1. For example, for the most simple implemented geometry contained in Ithildin, i.e., isotropic diffusion (see `Geometry_Iso` in sec. 4.1 and tbl. 2), the CFL condition is enforced by setting [21]:

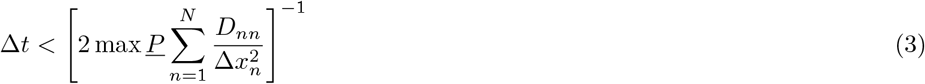

where *D*_*nn*_ are the diagonal components of ***D*** and max *<u>P</u>* is the maximum value of *<u>P</u>* .

**Table 2.** Overview of tissue geometries supported by Ithildin.

| class | description | dim $N$ | points in stencil |
| --- | --- | --- | --- |
| **Geometry_Iso** | isotropic diffusion | 1..3 | $2N + 1$ |
| ↳ **Geometry_ND** | isotropic diffusion in higher dimensions [27] | $N \in \mathbb{N}$ | $2N + 1$ |
| **Geometry_OrtAniso** | orthotropic diffusion | 2..3 | $\begin{cases} 9 & N = 2 \\ 19 & N = 3 \end{cases}$ |
| **Surface** | 2D domain with extrinsic curvature | 2 | 9 |
| **QuadricSurface** | quadric surface with rotated parallel fibers | 2 | 9 |
| ↳ **Ellipsoid** | ellipsoidal surface with rotated parallel fibers | 2 | 9 |
| ↳ **Hyperboloid** | hyperboloidal surface with rotated parallel fibers | 2 | 9 |
| ↳ **Paraboloid** | paraboloidal surface with rotated parallel fibers [28] | 2 | 9 |

Higher accuracy at the cost of more computations per time step, can be achieved with the RK4 method [19]:

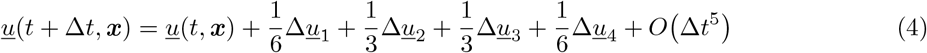

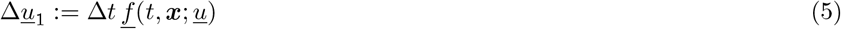

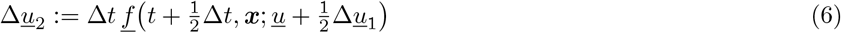

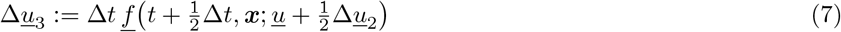

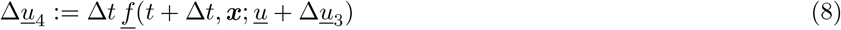

Note that *<u>f</u>* needs to be evaluated four times for the RK4 method and only once for the Euler method. Due to the higher accuracy though, the stability of this method is better than the one of forward Euler, since the latter is limited by the CFL condition. Indeed, using RK4, stability of the simulation is ensured for larger Δ*t* compared to what the CFL condition would suggest.

The stepping scheme to be used can be chosen in Ithildin on a per-variable level via `Model::steppings.`

### 2.3 Numerical spatial derivatives

For the numerical solution of the reaction-diffusion equation (eq. 1), the spatial derivative in its right hand side must be computed, i.e., the diffusion operator *<u>P</u>* ∇ · ***D***∇ *<u>u</u>*. This is implemented as weighted sums of the value of *<u>u</u>* at neighboring vertices on the grid of the discretized domain. The weights for the calculation of the stencil depend on the chosen type of diffusion (sec. 4.1). The two main types in this software are a first order stencil, including only the nearest neighbors, and a second order stencil, including also the next to nearest neighbors.

In the simplest case (see `Geometry_Iso` in sec. 4.1 and tbl. 2), we consider isotropic and homogeneous diffusivity. The diffusion operator can then be computed via the Laplacian operator ∇^2^. This is done with a 5-point stencil for the 2D case, a 9-point stencil for the 3D case, etc., see also fig. 2, panel (a). Consequently, the weights are calculated as:

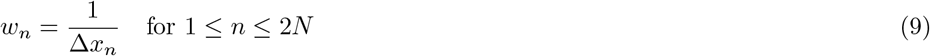

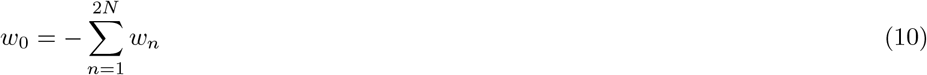

where *N* is the number of dimensions and Δ*x*_*n*_ is the grid resolution in the direction of the neighbor corresponding to weight *w*_*i*_, e.g., Δ*x*_5_ = Δ*z*. Note that the indices correspond to those displayed in fig. 2.

**Figure 2.**
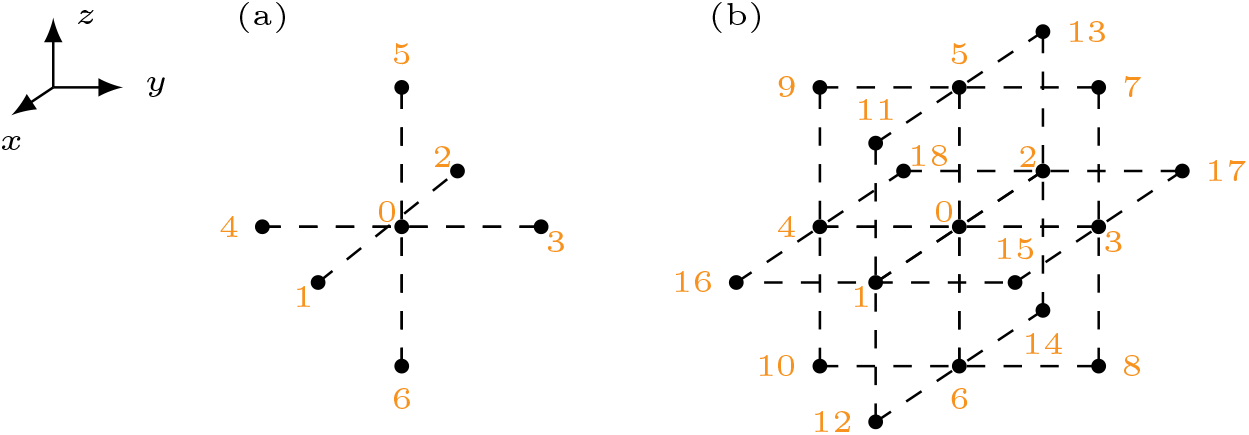
Stencils for isotropic diffusion (a) and orthotropic diffusion (b) with numbering of the involved grid points.

For the more general orthotropic diffusion, the stencil for numerical differentiation includes the nearest and diagonal neighbors of a grid point, see fig. 2, panel (b). The weights for orthotropic diffusion are obtained by a combination of central differences approximations to derivatives and linear interpolation of values between two grid points. For more details, the reader is referred to the documentation of `Geometry_OrtAniso.`

## 3 Implementation

The source code of Ithildin is written in the C++ programming language, following the 2011 version of the ISO-C++ standard [5]. This was chosen to facilitate programming at a level relatively close to the hardware, but also using some of the useful data-structures that are contained in the standard template library.

Ithildin is designed to run in parallel on multiple CPU cores using MPI, specifically OpenMPI [10]. Using the `mpirun` command, multiple instances, also known as processes, of the same compiled executable are started that run across different processor cores. The processes are then coordinated such that there is one so-called *manager* process, that manages a bunch of so-called *worker* processes. The manager does an equal share of the computational work, just like a worker. The only difference is that the manager distributes and directs information to be exchanged from one process to another. In Ithildin, the memory is not shared across processes, instead each process works on its share of the computational domain. We split the domain in the *x*-direction, such that each process is responsible for computations on a roughly equal share of vertices inside the to-be-simulated medium. Even when some parts of the domain are classified as exterior points, i.e., points on which no calculations need to be performed, they are taken into account when splitting the domain between processes. This splitting is possible because the reaction-diffusion systems to be studied with Ithildin are local, meaning that the temporal evolution at each time *t* and each point ***x*** in space depends only on the current state vector *<u>u</u>*(*t*, ***x***) at that position and its spatial derivatives (eq. 1). Internally, a layer of so-called *ghost points* is added around each process’ part of the domain such that the spatial derivatives can still be calculated in the same way as for any other point in the domain (sec. 2.3). The values *<u>u</u>* on these ghost points are exchanged with the neighboring processes, as coordinated by the manager process. Additional ghost points are also used to enforce Neumann boundary conditions, which is done by setting the weights for the calculation of the numerical spatial derivatives accordingly.

To run a simulation in Ithildin, the user needs to define a `main()` function that is to be called by all subprocesses. This is usually done using a C++ file calling the required components of the Ithildin library defining the `main()` function, a so-called *main file*. Ithildin can be installed as a shared library on Unix-based systems that is required by executables obtained from compiling main files. Alternatively, it is also possible to statically compile the Ithildin library with a main file into a stand-alone executable.

Besides the necessary preparations for using MPI, running a simulation using Ithildin’s Sim class requires three main components, which are instances of three classes:

1. `Model:` the reaction term *<u>r</u>*(*<u>u</u>*), typically a tissue model,
2. `Geometry:` the discretized diffusion term *<u>P</u>*∇ · ***D***∇ *<u>u</u>* for a chosen geometry of the medium, and
3. `Source:` the stimulus protocol to use as well as inhomogeneities in the medium.

In the following sec. 4, an overview is given for each of these classes and their derived classes. Combining all of the components, an illustrative, minimal main file can be obtained, in which a planar wave crosses the medium in positive *x*-direction:

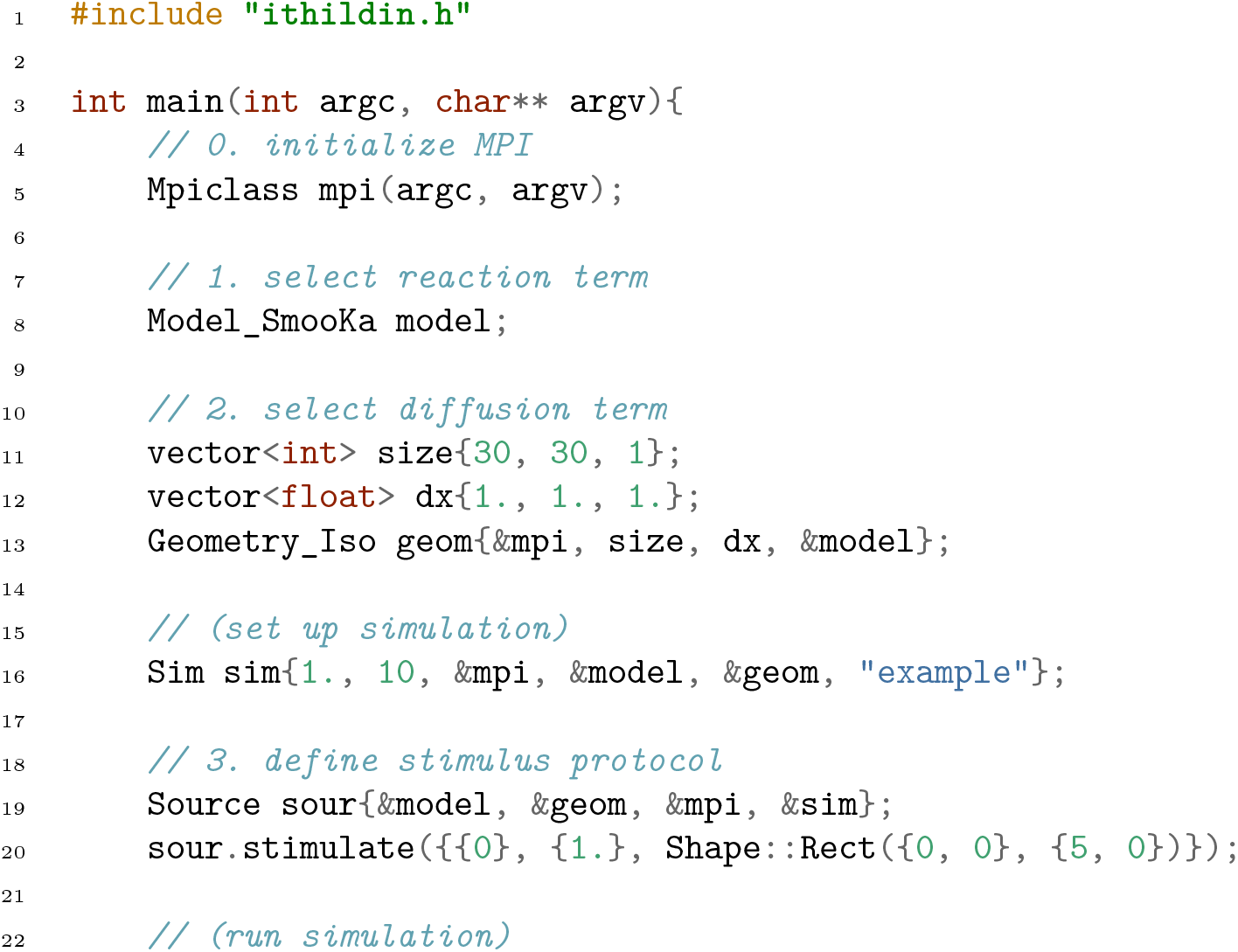
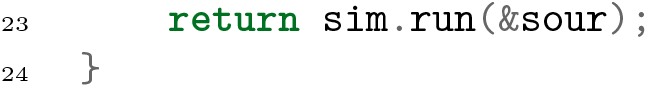

More detailed examples for main files can be found in the appendix, where we set up the numerical examples used throughout this paper.

We consider the geometry, the model and the source to be the inputs of Ithildin, see also fig. 1. Upon running the simulation, Ithildin produces a variety of outputs, as files in easy-to-parse standardized data formats. The names of these files all begin with the so-called *stem*, consisting of a descriptive series name and a serial number, which defaults to a time-stamp. In the following, an overview of the usual output files of Ithildin by suffix appended to the stem is provided:

- `_log.yaml:` The log file contains metadata describing the setup of the simulation, as well as metadata about the conditions under which the simulation was run. This file is always output by Ithildin and is considered the central file of the results, as it points to the relevant other files that are only conditionally written during the simulation. While earlier versions of Ithildin used a non-standard format for the log files, in current versions, the YAML format is used, making it easy to parse by both: humans and machines [12].
- `_main.cpp:` A copy of the C++ code in the main file may be included in the results for reproducibility.
- `_git.diff:` If run in a Git-repository with changes since the last commit, a patch file of these changes will be included in the results.
- `_*.txyz.npy:` For each of the state variables *<u>u</u>*, a so-called var file in the NumPy NPY format will be written [13]. These files contain the *N* + 1-dimensional floating-point number array of the evolution of a state variable *u* in the whole computational grid over time. Note that the order of indices is (*t, x, y, z*), meaning that time is the slowest varying index and the *x*-axis the second slowest varying index. This order was chosen because the domain is split across processes along the *x*-axis, such that the processors can open and write to these files sequentially. For 2-dimensional simulations, the third spatial dimension, the *z*-axis, is one vertex thick, such that each var file still has four dimensions. For higher-dimensional simulations, more axes are added, leading to more dimensions in the var files.
- `_inhom.txyz.npy:` The inhom field, see details in sec. 4.3, is also stored in the NPY format, but with integer values, and only for the initial time-step. This file hence has the shape (1, *N*_*x*_, *N*_*y*_, *N*_*z*_) with *N*_*n*_ denoting the number of vertices in each of the spatial dimensions.
- `_hist*.csv:` Comma-separated value (CSV) files describing the temporal evolution of *<u>u</u>*(*t*, ***x***_s_) at a chosen sensor position ***x***_s_ ∈♡ ℝ⊂^*N*^, see details in sec. 4.5.
- `_egm*.csv:` CSV files containing the recorded pseudo-EGM Φ(*t*, ***x***_e_) at a chosen electrode location ***x***_e_ ∈ ℝ^*N*^, see details in sec. 4.6.
- `_tipdata.yaml:` This YAML file is written if filament-tracking is turned on, see also sec. 4.7, and contains the detected phase singularities in regular time intervals.

While we have selected these file formats to be easy to read using a wide variety of software, we have also developed the Python module for Ithildin to facilitate interacting with the results of an Ithildin simulation and converting them to a variety of file formats, for instance writing files in the extensible data model and format (XDMF) that can be used to view simulation results in ParaView [14, 17, 22, 23]. The Python module also offers post-processing and analysis methods, for instance the computation of action potential duration (APD), conduction velocity (CV), various phases, phase defect detection, functions acting on filaments and filament trajectories, and several plotting functions [14, 22].

## 4 Application-focused numerical methods

### 4.1 Diffusion term

The diffusion term in the reaction-diffusion equation (eq. 1) is stated as *<u>P</u>*∇ · ***D***∇*<u>u</u>*. The conduction in cardiac tissue and hence the diffusion is stronger along the fiber direction than normal to the fibers [2]. This is encoded in the diffusion matrix ***D***.

As an example, in fig. 3, the fiber direction is drawn on the surface of the ventricular geometry used in sim. 4. The fibers are additionally colored by their fiber helix angle [24]. The voxel-based representation of this geometry was obtained by cutting a human heart into 1 mm-thin slices, digitizing and stacking them [25, 26].

**Figure 3.**
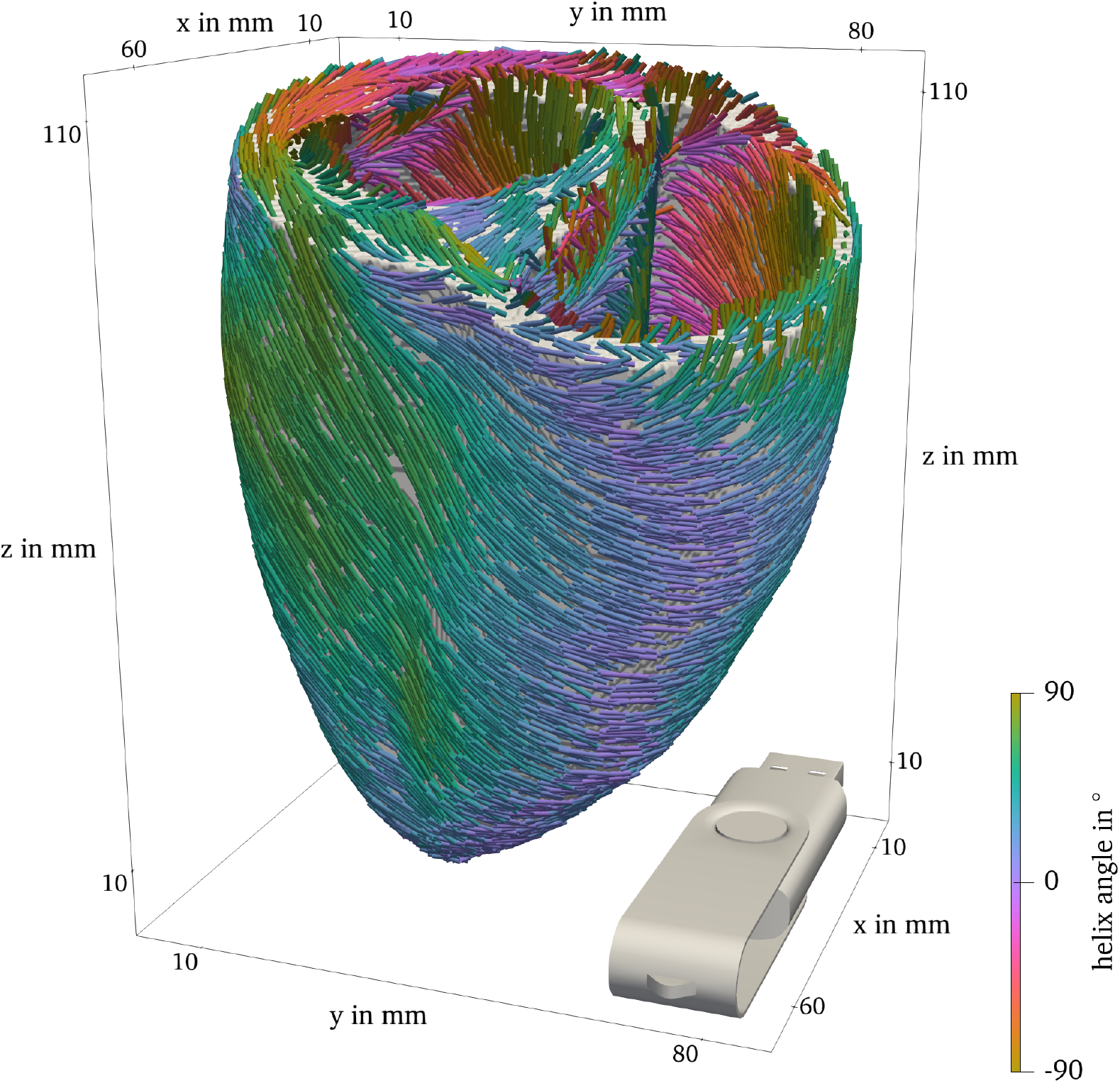
Ventricle geometry with fiber direction, colored by the fiber helix angle. USB flash drive for scale and orientation. (3D model of a USB flash drive by Dmytro Rohovyi, licensed under CC-BY-NC.)

While the selection matrix *<u>P</u>* is managed by the Model class, see sec. 4.2, the diffusion matrix ***D*** and the handling of the spatial derivatives is implemented in the `Geometry` class. The most important things the `Geometry` class takes care of, are:

- Initialization of the computational grid, along with the strides and pointers, which are important for efficient computing;
- Computation of the entries of the diffusion tensor, based on the main directions of diffusion and the respective diffusion values;
- Computation and storage of the weights for the stencils of the numerical spatial derivatives (sec. 2.3) respecting the Neumann boundary conditions; and
- Handling the upper bound for the time step due to the CFL condition (sec. 2.2).

The `Geometry` class is a base class and should not be used directly to construct a computational domain. Instead, there are several subclasses, each representing a different type of diffusion or extrinsic shape. An overview of the `Geometry` subclasses is given in tbl. 2.

The `Geometry_ND` class is a subclass of `Geometry_Iso` and the `Ellipsoid, Hyperboloid,` and Paraboloid classes are subclasses of `QuadricSurface.`

The details on how the extrinsic and intrinsic curvature affect the diffusion tensor and hence the weights for the discretized differential operator, are discussed in the documentation of the code. There, the different ways to initialize the available domain types are given as well.

Note that some additional features are implemented in the `Geometry` class, that do not have a direct link with the diffusion term, such as inhomogeneities (sec. 4.3) and filament detection (sec. 4.7).

### 4.2 Reaction term

The Model class serves as the base class for all cardiac electrophysiology models in the Ithildin framework. It encapsulates common functions and variables used across different models. Key features of the `Model` class include:

- Implementation of the reaction term *<u>r</u>*(*<u>u</u>*) in the reaction-diffusion equation.
- Storage of model-specific metadata such as relevant citations.
- Handling of variable-related information, including their names, indices, and resting values.
- Management of projection matrix *<u>P</u>* values for variables, which is typically a diagonal matrix describing whether or not a variable is diffused. For instance, for the AP96 model [29], *<u>P</u>* = diag (1, 0).

Derived classes extend the `Model` class to implement specific cardiac electrophysiology models in the `reactionterm` function.

An overview of the tissue models that are currently included in the source code of this project is provided in tbl. 3. More models may be added as additional classes.

**Table 3.**
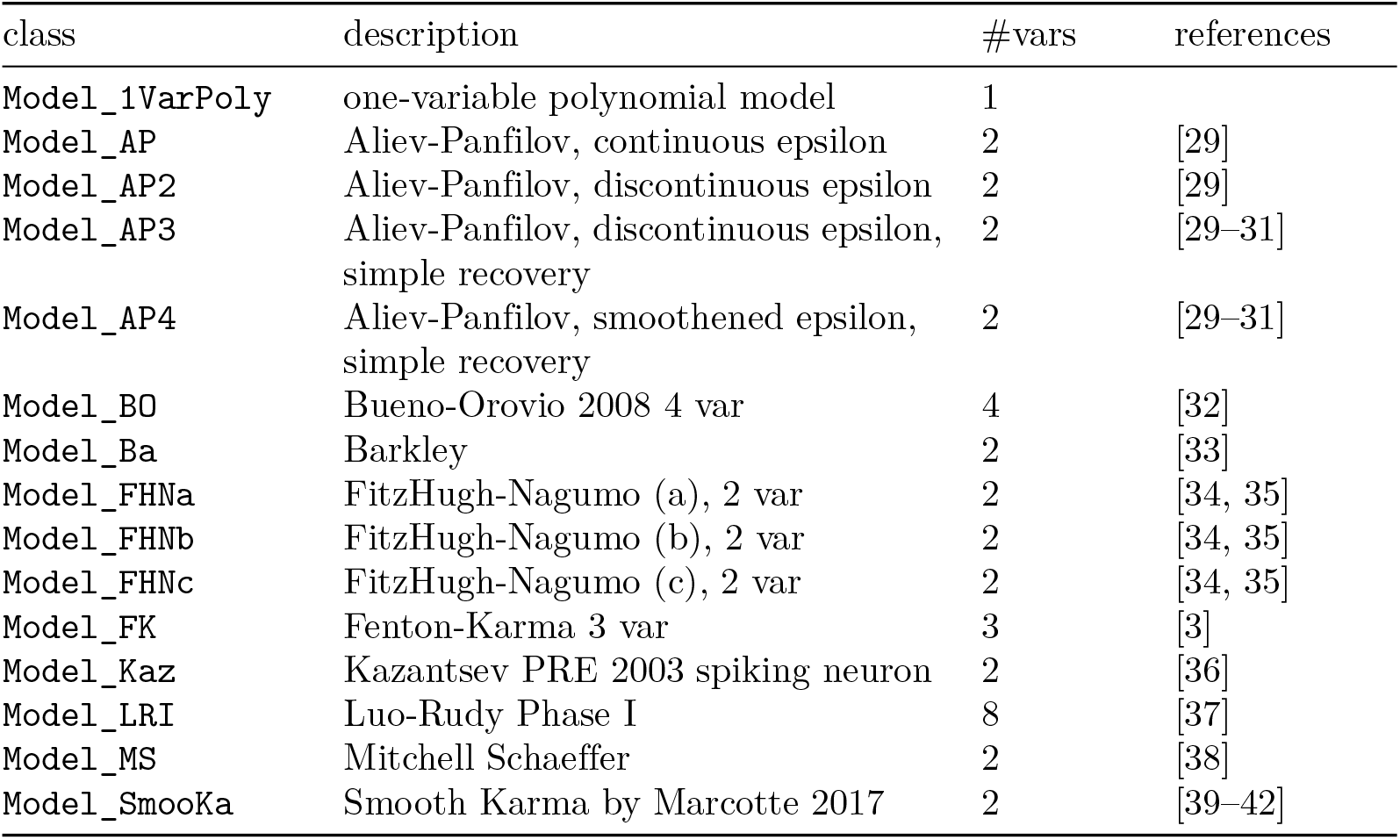
Overview of tissue models included in Ithildin.

The Ithildin framework also introduces several `ModelWrapper` classes that provide additional functionality and allow for the combination of models. These wrappers enable the recording of diffusion terms, reaction terms, local activation times (LAT), and local deactivation times (LDT) as additional state variables. This is done by adding code to the reaction term calculations of the underlying model. The wrappers inherit from the base `ModelWrapper` class, which is a wrapper that leaves the model unchanged. The primary `ModelWrapper` classes are:

- `ModelWrapper_RecordDiffusion` records diffusion terms of selected variables as additional variables.
- `ModelWrapper_RecordReaction` records reaction terms of selected variables as additional variables.
- `ModelWrapper_RecordActivationTime` records LAT for selected variables.
- `ModelWrapper_RecordDeactivationTime` records LDT for selected variables.
- `ModelWrapper_RescaleTimeSpace` linearly rescales the wrapped model in time and space.
- `ModelWrapper_RescaleVars` linearly rescales selected state variables of the wrapped model.

The `Model_multi` class enables the combination of multiple submodels into a single model. The behavior of the combined model may vary depending on the location ***x*** and is determined by one of its submodels. Which model is to be used depends on the integer value of the `inhom` field, which is used to describe spatial inhomogeneities, such as obstacles. Inhomogeneities will be further explained in the following, cf. sec. 4.3. This class is particularly useful for simulating scenarios where different regions of cardiac tissue exhibit distinct behaviors.

The Ithildin C++ framework provides a structured and modular approach to modeling cardiac electrophysiology. The `Model` class serves as the base for different models, while various `ModelWrapper` classes and the `Model_multi` class offer extended functionalities for recording and combining different model aspects.

### 4.3 Inhomogeneities

In simulations of cardiac electrophysiology, accurately modeling the spatial properties of the cardiac tissue is essential. Inhomogeneities represent variations in the tissue’s characteristics, such as its electrical conductivity or cellular properties, that influence the propagation of electrical signals.

An inhomogeneity in the Ithildin framework is a distinct region within the simulation domain with different properties compared to its surroundings. In the context of cardiac electrophysiology, these properties could correspond to variations in the electrical conductivities of cells, cellular properties, or even the absence of excitable cells altogether. Inhomogeneities are defined by an integer field called `inhom` associated with each point in the domain. Grid points with a non-zero `inhom` value are considered *interior points*, indicating that the reaction-diffusion equation needs to be solved on these points. If the selected reaction term is a `Model_multi` (see also sec. 4.2), for an `inhom` value of *n*, the *n*th submodel will be used for this point. For example, in fig. 4, the `inhom` field for sim. 1 is visualized. Two different tissue models are used for the values 1 and 2. The points where `inhom` has the value 0, are considered *exterior points*. The set of all interior points is the physical domain ♡. At the boundary of the physical domain, Neumann boundary conditions are applied.

**Figure 4.**
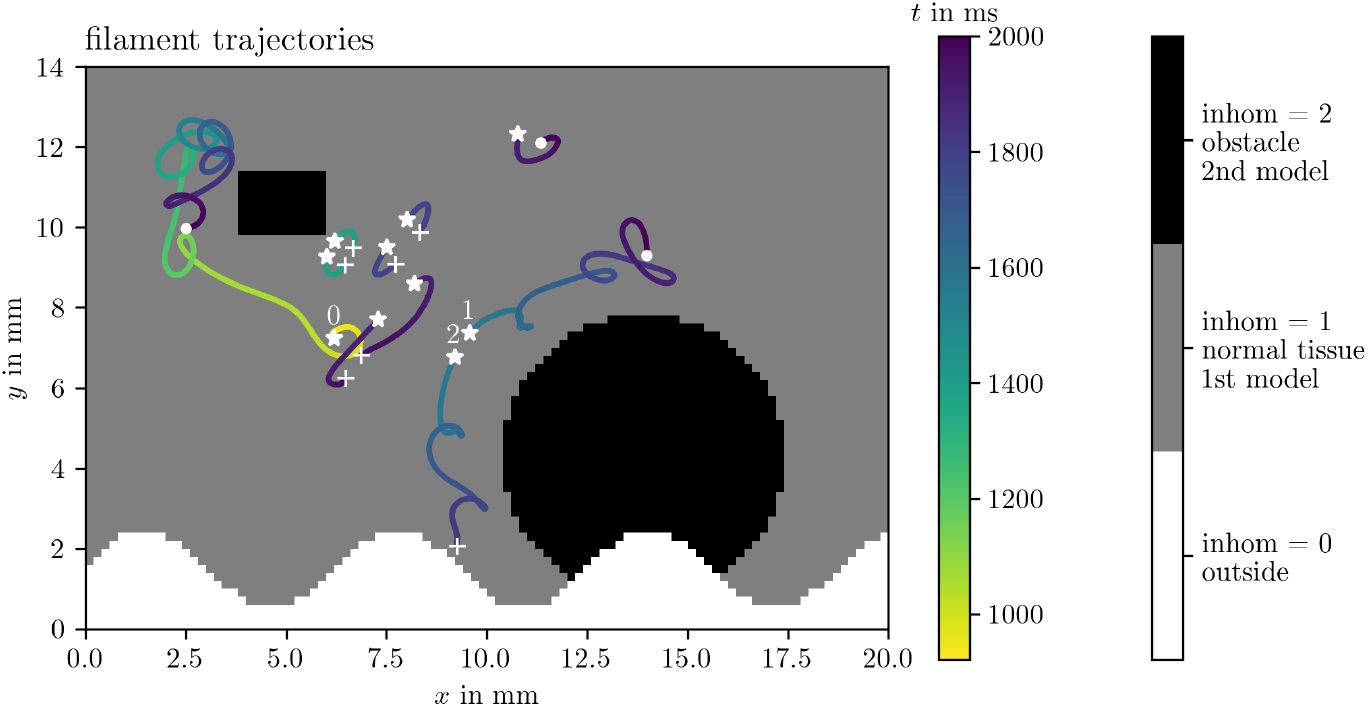
The field `inhom`, shown here for sim. 1, describes where unexcitable obstacles or exterior points are located with the value `inhom` = 0 and which tissue model is to be used inside if `inhom > 0`. This figure also visualizes the filament trajectories colored by time, cf. sec. 4.7. Birth of a tip is denoted by stars, their deaths by crosses, and tips that are still around at the end of the simulation by points. The three trajectories with the longest lifetime are numbered by indices 0 through 2.

To incorporate inhomogeneities into the simulation, the framework provides methods to add them to the domain. These methods allow specifying the shape, location, and properties of each inhomogeneity. For example, a rectangular inhomogeneity could be added by specifying its width, height and location.

### 4.4 Stimulation protocols

Ithildin provides a flexible way to define stimulation protocols using the `Stimulus` and `Trigger` classes, as well as the scheduling functionality of the `Source` class.

The `Stimulus` class enables the specification of temporal and spatial characteristics of voltage-based or current-based stimuli. It allows defining which state variables should be affected directly, the associated values, and whether the stimulus sets the variable directly or whether it is additive and hence behaving like a current source. Temporal modulation of stimulus strength is achieved through the `amplitude` function, while spatial constraints are managed by the `shape` function, an instance of the `Shape` class.

The `Shape` class describes a geometric shape via its characteristic function which can be used to define regions of interest in a simulation domain. The core principle is that the function evaluates a given position vector and returns a value: 1 if the position is inside the defined shape, and 0 if it is outside. However, values in the range [0, 1] may also be used to create a smooth transition. A smoothly varying characteristic function may be useful to create more-realistic stimuli that deposit current in a smooth profile. This simple yet powerful concept forms the basis for constructing intricate spatial configurations.

The Shape class provides several pre-defined shape functions, though additional shapes can easily be added by defining a characteristic function:

- **Ellipsoid**: Defined by radii, a center and optionally the Euler angles, this shape represents a general three-dimensional ellipsoid with the specified orientation.
- **Sphere**: A special case of an ellipsoid where all radii are equal, forming a three-dimensional sphere.
- **Ellipse in** *xy***-plane**: This two-dimensional shape resembles an ellipse lying on the *xy*-plane, defined by radii and a center.
- **Cylinder along** *z***-axis**: Representing a three-dimensional cylinder centered along the *z*-axis, this shape is defined by a radius and a center. In 2D, it defines a disk.
- **Rectangular cuboid**: Defining a three-dimensional region, this shape is specified by two opposing corner points, creating a cuboid. In 2D, it defines a rectangle.
- **Half plane**: A plane defined by an origin and an outward normal vector splits the three-dimensional space into a half-space. In 2D, a straight line splits the plane in a similar way.

Characteristic functions can also be loaded from files in the NPY format [13]. This feature facilitates the incorporation of custom shapes derived from external data sources.

The scheduling functionality via `Source::schedule` offers the ability to execute functions at specified points in time during the simulations. This feature greatly enhances experimental flexibility by allowing the execution of arbitrary code snippets at chosen moments during simulations. The scheduler is especially useful for introducing dynamic changes to the simulation environment, such as modifying stimuli or conditions mid-simulation.

The `Trigger` class provides a means to orchestrate actions based on specific conditions. Triggers encapsulate the decision-making process of when to execute a particular action, influenced by condition checks and coordination modes. Different coordination modes allow for the synchronization of trigger actions across multiple processes, facilitating complex simulations of activation waves. These modes are to trigger on each process individually once the condition is met, once the condition is met in a specific process, once the condition is met in any process, or once it is true in all processes.

Within the framework of cardiac electrophysiology, triggers are essential for defining stimulation protocols, e.g. for the S1S2 protocol, which is illustrated in fig. 5: After a first excitation wave passes a sensor position, a second wave is triggered behind a part of the first waveback to stimulate spiral waves.

**Figure 5.**
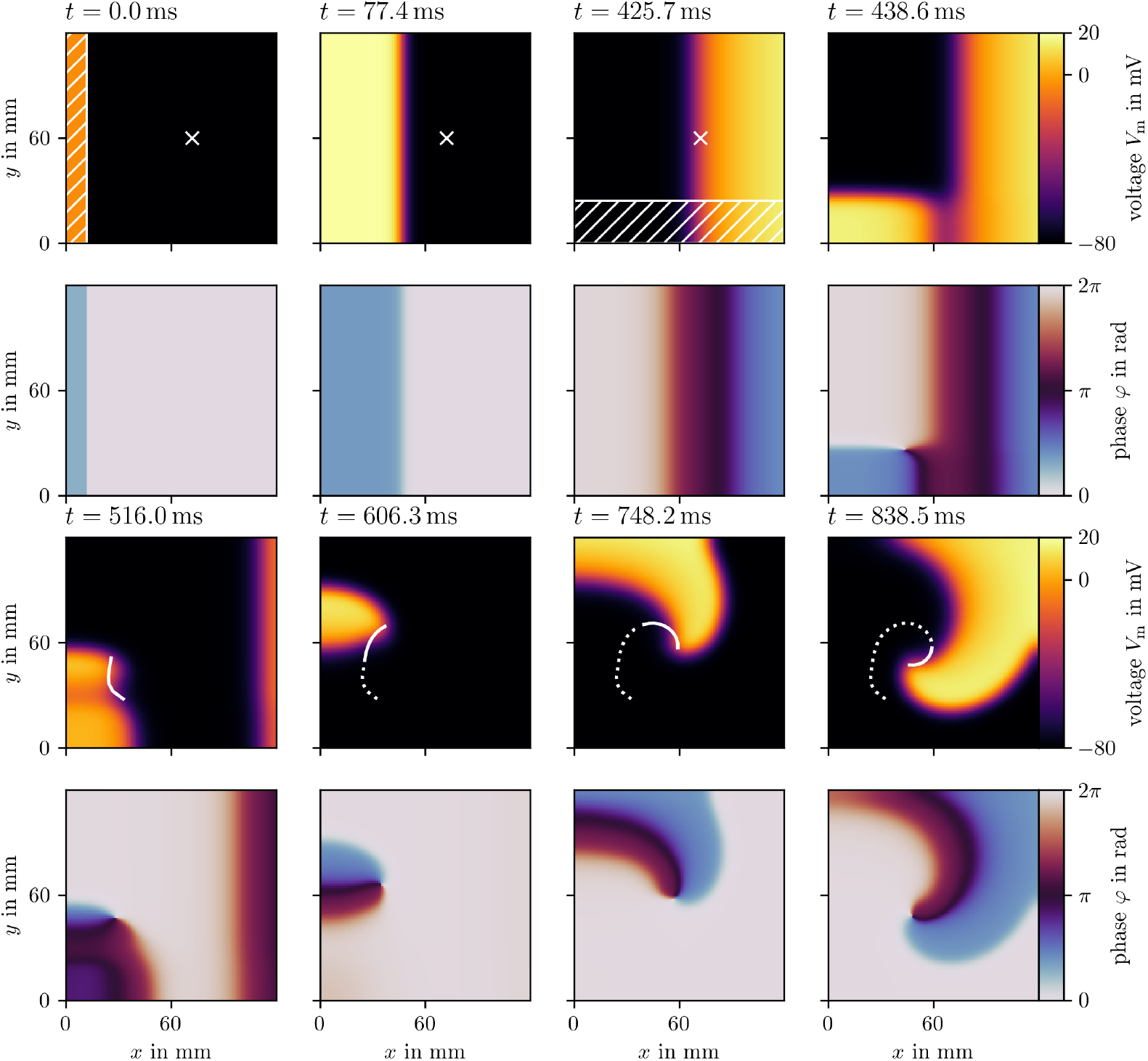
The S1S2 protocol illustrated for sim. 2. The first and third row display the transmembrane voltage *u* at selected frames in time, and the second and fourth row the state space phase [14, 43]. The first stimulus is applied in the first frame in the hatched region at the left edge. The sensor location is marked with a cross. Right after the third frame, the sensor triggers the second stimulus in the hatched region at the bottom edge. A spiral wave forms. The phase singularity at the center of the spiral is tracked as the white curve, which is dotted for the entire trajectory of the phase singularity, and solid for its trajectory since the previous frame.

### 4.5 Recording temporal evolution of variables at sensor positions

Besides the sensors for triggering stimuli, sensors in the context of this simulation framework are components that monitor and record the state variables of the simulated system at chosen positions during the simulation. We call this the *history* at a given sensor position.

These sensors are used to gather data about the behavior of the system at particular time intervals, regulated by the `sensorlag` parameter. This parameter controls the frequency at which sensor data is collected, allowing for flexibility in recording intervals. Notably, the recording frequency set by `sensorlag` need not align with the simulation’s time step or the duration between frames. The data will be recorded at the first time step after the specified `sensorlag` duration. This high-resolution temporal data can be used to study individual points in the medium in detail.

The recorded data are then written to designated comma separated value (CSV) output files associated with each sensor. These files are used to store the collected data over the course of the simulation.

The first four panels of fig. 6 contain time traces of the transmembrane voltage *u*, the restitution variable *v*, the recorded value of the diffusion term, and the LAT at a given sensor position for sim. 1. The times of the three stimuli are indicated by the black vertical lines. The other panels will be explained in the subsequent sections.

**Figure 6.**
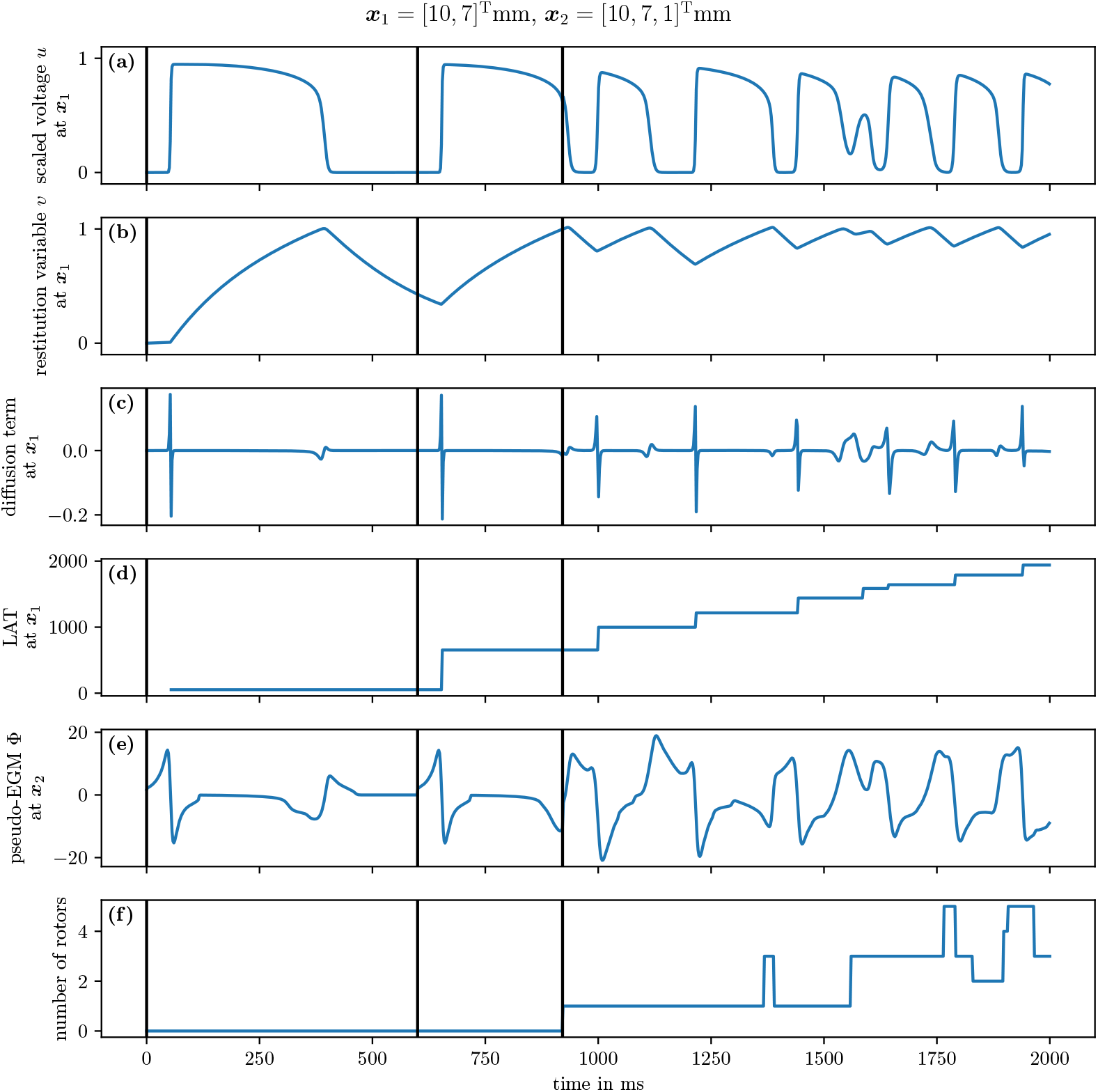
Extracted time traces of sim. 1. The first two panels (a, b) contain the model variables *u* and *v*, followed by data computed by model wrappers, namely the diffusion term∇ · ***D***∇*u* (c) and the LAT (d), at an interior point, cf. sec. 4.5. Panel (e) contains the pseudo-EGM at an exterior point, next to the 2D domain, cf. sec. 4.6. The last panel (f) contains the number of rotors detected via phase singularities, cf. sec. 4.7. The black vertical lines indicate times at which a stimulus was applied.

### 4.6 Pseudo-EGMs

The EGM is a measurement of the potential generated by the charge distribution in cardiac tissue over time at a point in space, outside the tissue. In theory, this is a measurement of the extracellular potential Φ_e_. However, since Ithildin is a mono-domain solver, the extracellular potential is not part of the model equations [2]. The code therefore calculates an approximation of the extracellular potential at points outside the mesh using the `Egm` class. To obtain this approximation, the pseudo-bidomain theory is used [44]. The approximation, referred to as a pseudo-EGM, uses the simplifications that the intracellular and extracellular conductivities are proportional, such that there is an explicit formula to calculate the extracellular potential, and that the bath conductivity is homogeneous.

The extracellular potential Φ_e_ at position ***x***_e_ over time is then calculated as [2]:

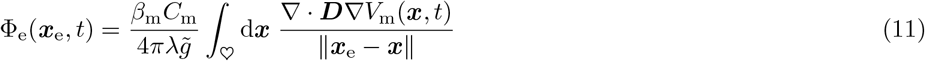

where ♡ denotes the computational domain, i.e., the simulated heart muscle tissue, *V*_m_ is the transmembrane potential, which is encoded in the state variables *<u>u</u>* of the model, usually as the first state variable *u*. ***D*** is the diffusion matrix from the reaction-diffusion equation (eq. 1), *β*_m_ is the surface to volume ratio of a cardiac cell, *C*_m_ is the specific cell membrane capacitance, *λ* is a proportionality factor between intracellular ***G***_i_ and extracellular conduction matrix ***G***_e_, such that ***G***_e_ = *λ****G***_i_, and 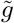 is a lumped conductivity.

Essentially, the used approximation for the EGM is a convolution of the diffusion term ∇ · ***D***∇*V*_m_ for the first variable with a kernel ∥***x***_e_− ***x***∥ ^−1^. Depending on the choice of the diffusion tensor, made by the user in the `Geometry` class (sec. 4.1), a different prefactor of the integral is required. Hence, the prefactor is user-defined and can be changed using the function `set_prefactor.`

The integral in eq. 11 is discretized and its calculation is implemented such that the additional amount of storage and number of calculations is limited as much as possible. For instance, as the required diffusion term ∇ · ***D***∇*V*_m_ is already computed during the forward stepping of the reaction-diffusion equation, it can be stored as an additional state variable using a `ModelWrapper_RecordDiffusion` and subsequently used in the pseudo-EGM calculation. This model wrapper is essential for the functioning of the `Egm` class and hence is a requirement when setting up a simulation with pseudo-EGM calculation.

The result is a CSV file with the pseudo-EGM data for each electrode that is defined by the user. The computed pseudo-EGM at a given position for sim. 1 is displayed in the fifth panel of fig. 6.

### 4.7 Filaments

Formally, filaments can be understood as a line of wave break, i.e., a line where an activation and recovery surface come together [45, 46]. The activation surface can be seen as the wavefront, while the recovery surface can be seen as the waveback. When considering an excitable system in 2D, filaments become tips, being the point of intersection between the activation and recovery curve. Since a point cannot be excited and recovering at the same time, points on a filament are also called *phase singularities*.

Using this definition of filament points, detection algorithms have been designed. Our code relies on the one described by Fenton *et al*. [3]. Additionally, this algorithm has been extended to grids in any dimension, where the generalization of filaments are called *superfilaments* [27].

The filament point detection algorithm is included in the `Geometry` class. Its goal is to compute the points where the wavefront and waveback meet. While looping over all coordinate planes and grid points, it is checked whether there is an intersection of isolines in the adjacent voxel faces of a grid point. The location of the filament point is then estimated by bi-linear interpolation. An illustrative sketch of this method is given in fig. 7.

**Figure 7.**
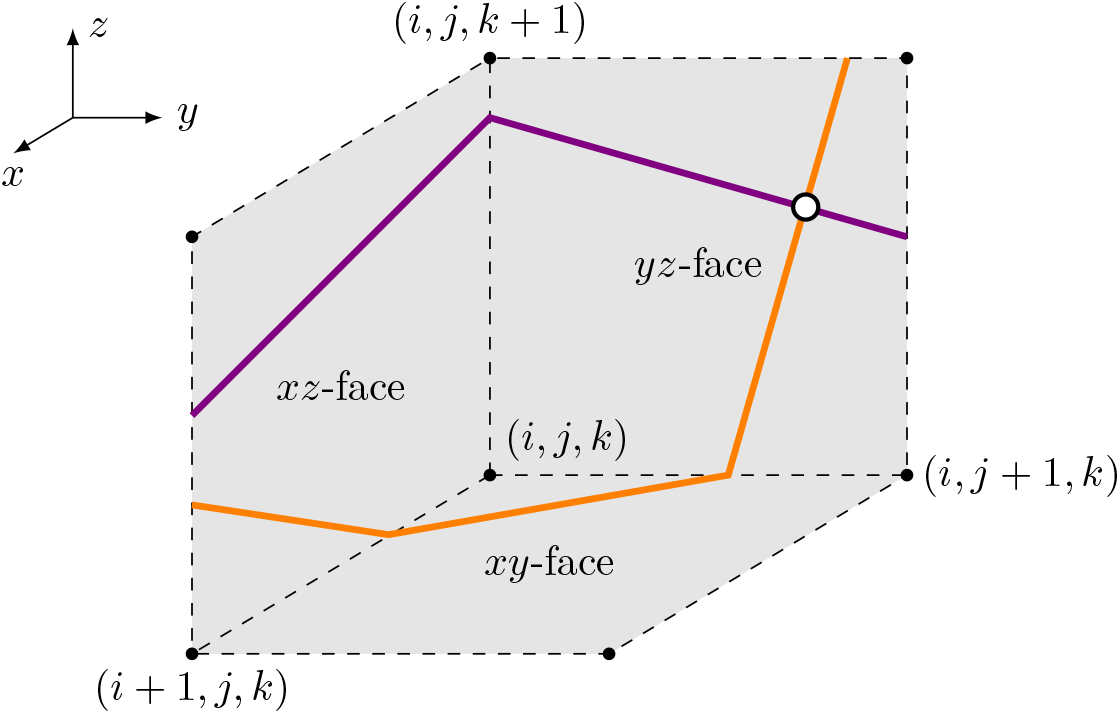
Illustration of the filament point tracking algorithm. Each coordinate plane adjacent to a grid point (*i, j, k*) is checked for an intersection of two surfaces (purple and orange lines) which are usually isosurfaces of state variables in *<u>u</u>*. In this example, a filament point (white dot) will be found in the *yz*-face.

The last panel in fig. 6 displays the number of rotors over time for sim. 1, which are found via filament detection. It can be seen that at the S2 stimulus, a single rotor is formed, and subsequently pairs of rotors as figure-of-eight spiral pairs.

More details of this process can be seen in fig. 4 which shows the trajectories of these filaments in sim. 1 tracked using the Python module for Ithildin. The trajectories are colored by time and the formation of a new tip is denoted by stars and their decay by crosses. Tips that still persist at the final frame of the simulation are indicated by points. The tip trajectory denoted with index 0 is formed by the S1S2 protocol and meanders around the medium. It persists until the end of the simulation. The figure-of-eight spiral wave pair denoted by indices 1 and 2 is formed by a conduction block breaking up. The spiral tip with index 2 runs into the boundary to disintegrate there.

### 4.8 Phase defects

While a phase singularity can be seen as points where all phases meet—in mathematics called a *pole*, a phase defect is a point at which there is a discrete jump from one phase value to another in an otherwise continuously varying phase. While phase defects are well known in physics, they were only recently identified in excitable media, within linear-core rotors and conduction block regions [43, 47]. Phase defects are lines in 2D and surfaces in 3D.

In Ithildin, the phase defect detection is done with its Python module. For instance, during simulation, Ithildin can record the local activation times (LAT) which can then be used to compute the activation time phase in post-processing [14, 43]. A variety of methods exist to then localize the phase defect [14, 47].

Two examples for phase defect detection in sim. 1 are given in fig. 8 and fig. 9. Both display four frames over time of the transmembrane voltage *V*_m_, the activation time phase *φ*, and the phase defect *ϱ* computed via the cosine method [14, 47]. In fig. 8, the phase defect of a single spiral wave is tracked. It can be seen that the phase defect extends due to conduction block such that the spiral wave moves across the domain. In fig. 9, the break-up of a conduction block line into a figure-of-eight spiral wave pair can be seen. The conduction block line is a phase defect of zero topological charge [48–50] which, in this case, reaches a critical length breaking apart into two oppositely charged spiral waves with much shorter phase defect lines.

**Figure 8.**
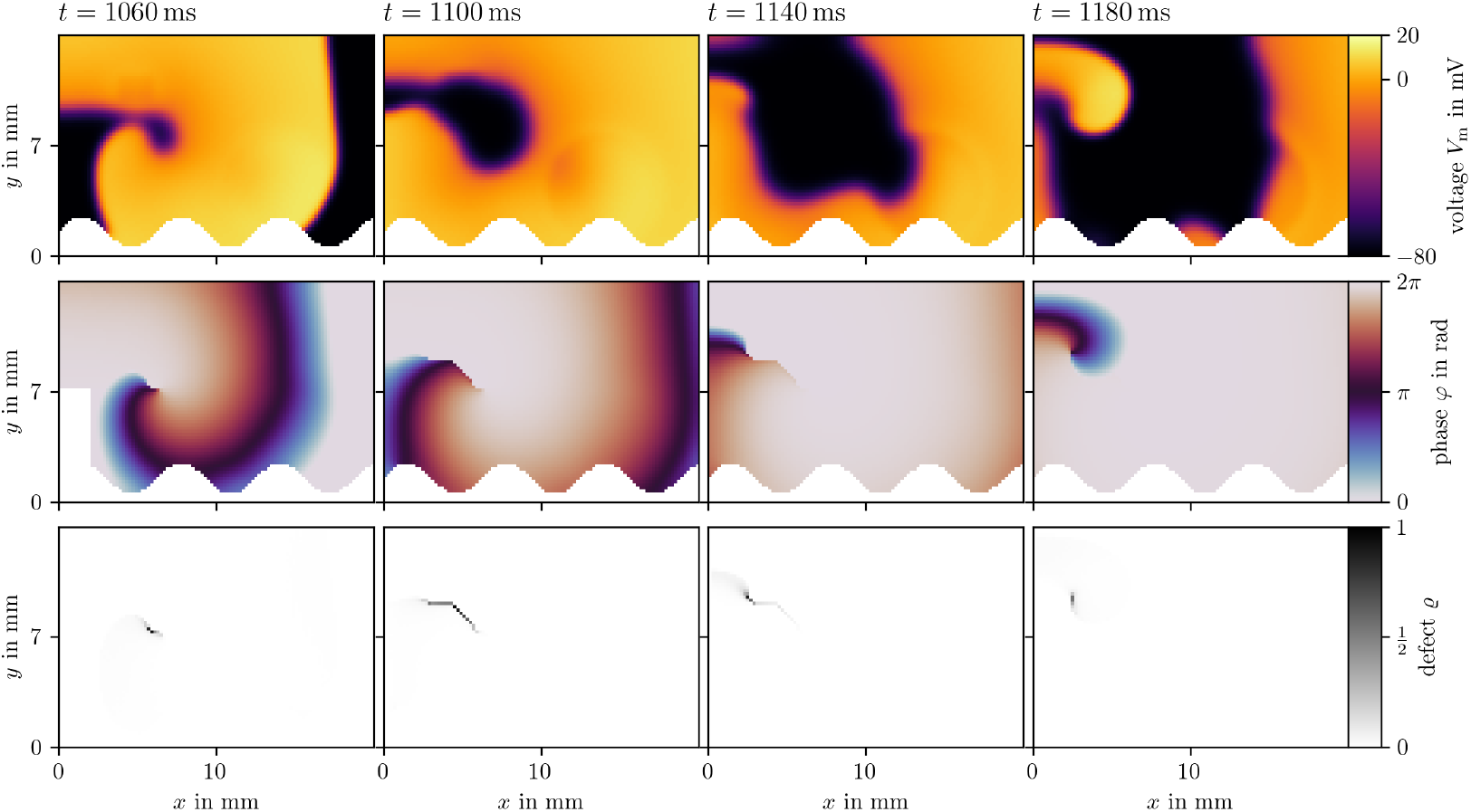
Phase defect detection for a single meandering spiral in sim. 1. By recording the activation times of the transmembrane voltage *V*_m_ (top row), the activation time phase *φ* [14, 47] can be computed (middle row). Where this phase is discontinuous, a phase defect is localized. This is visualized as the phase defect density *ϱ* (bottom row). In these four frames, a single rotor is tracked shortly after its formation. Due to conduction block, which can be seen as an extended phase defect line, the spiral moves through the medium. Afterwards, the spiral remains mostly stationary, leading to a shorter phase defect line.

**Figure 9.**
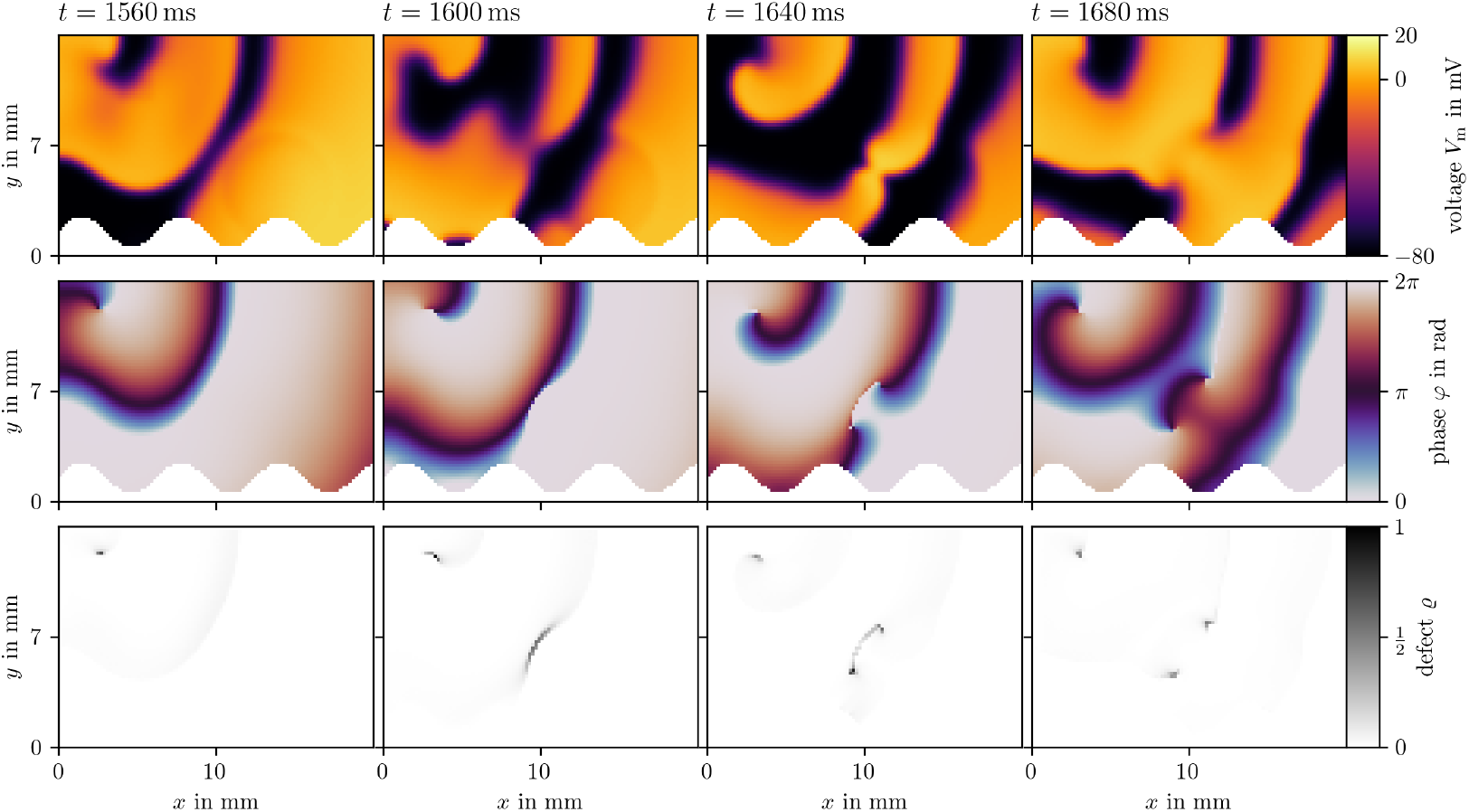
Phase defect detection during figure-of-eight spiral wave pair creation in sim. 1. Visualization in the same style as fig. 8. A long conduction block line breaks apart into two rotors.

In fig. 10, we present the final frame of sim. 4 in ventricular geometry, visualized with ParaView [17]. It is colored by the normalized transmembrane voltage. Both, the classical tip and the phase defect surface are visualized. The spiral waves revolve around the phase defect surfaces.

**Figure 10.**
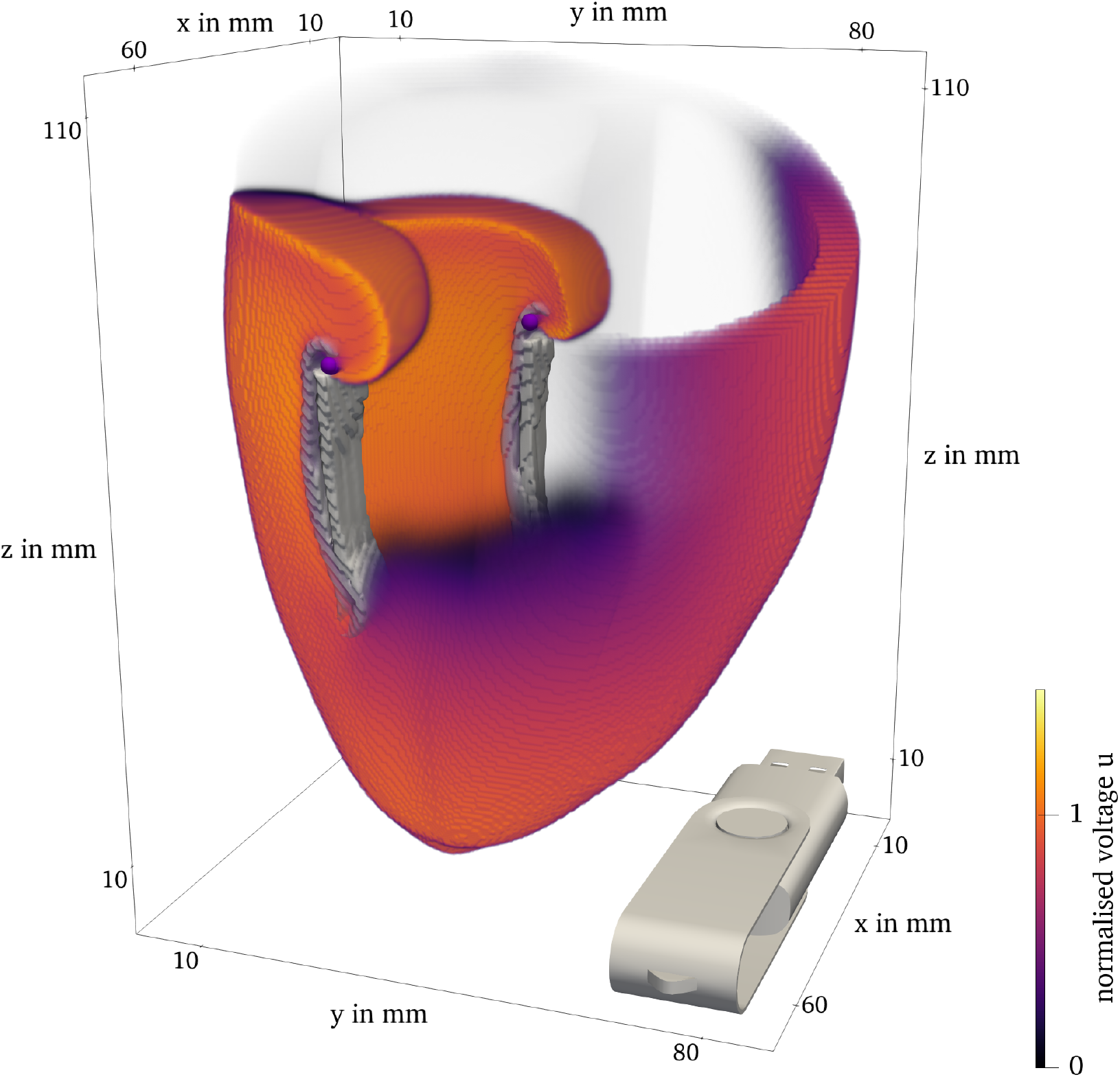
Final frame of sim. 4 of the BOCF model in ventricle geometry. The classical filament is plotted as the purple lines. The phase defect surface is contained in the gray contour. USB flash drive for scale and orientation. (3D model of a USB flash drive by Dmytro Rohovyi, licensed under CC-BY-NC.)

### 4.9 Further documentation

More complete documentation of Ithildin can be generated with Doxygen [11] and can be also found online, see the data availability statement for details. The documentation contains detailed information on how to get started installing and working with Ithildin.

## 5 Results

Ithildin is used for numerical experiments in various use cases. For instance, it was used to study the structure of the core of rotors emerging in in-silico cardiac-electrophysiology tissue models, leading to the description of phase defect lines [14, 43, 50, 51]. Also, higher-dimensional rotors waves were simulated and the emerging super-filaments were detected using Ithildin [27]. In the creation of novel data-driven tissue models using state space expansion, Ithildin was used to generate synthetic training data sets [22].

Four simulations were conducted for this paper to illustrate the features of Ithildin. An overview of the simulations is given in tbl. 4, along with references to the figures that were generated with these data sets while details on their simulation setup can be found in their C++ code in the appendix.

**Table 4.** Overview of the numerical simulations used in this paper.

| simulation | sim. 1 | sim. 2 | sim. 3 | sim. 4 |
| --- | --- | --- | --- | --- |
| tissue model | Model_SmooKa | Model_AP | Model_BO | Model_BO |
| parameter set | default [39–42] | default [29] | EPI [32] | EPI [32] |
| dimensions | 2 | 2 | 2 | 3 |
| anisotropy type | isotropic | isotropic | isotropic | orthotropic anisotropy |
| diffusivity $P_{uu}$ | 0.031 | 1.6 | 0.12 | 0.12 |
| fiber ratio $D_{\parallel}/D_{\perp}$ | 1 | 1 | 1 | 4 |
| inhomogeneities | sphere, rectangle, waves | none | none | biventricular geometry |
| grid size | $100 \times 70$ | $120 \times 120$ | $450 \times 450$ | $168 \times 208 \times 231$ |
| resolution [mm] | 0.2, 0.2 | 1, 1 | 0.3, 0.3 | 0.43, 0.43, 0.5 |
| time step $\Delta t$ [ms] | 0.1 | 0.1 | 0.1 | 0.1 |
| source times [ms] | 0, 600, 921 | 0, 426 | 0, 394.1 | 0 |
| duration [ms] | 2000.1 | 838.6 | 680.1 | 600.1 |
| used in | figs. 1, 4, 6, 8, 9 | fig. 5 | sim. 4 | figs. 1, 3, 10 |

## 6 Discussion

The Ithildin framework is a tool for the simulation of cardiac electrophysiology. The code offers a number of assets that allow for simulations targeting a variety of phenomena. For example, there are many instances available to set the local anisotropy of the myocardium via the `Geometry` class. Please refer to the example presented in sec. 4.1 for further details. The `Geometry` class also allows for the simulation and filament tracking in a domain with an arbitrary number of spatial dimensions [14, 27, 43, 50, 51].

These include the ability to define inhomogeneities, which can be used to model domains of any shape and with any kind of obstacles. This is used in sim. 1 and sim. 4. Furthermore, the user can define a variety of stimulation protocols, of which some are showcased in the example simulations. We also note the following options: EGM calculation, the possibility of recording the temporal evolution of variables at sensor positions, and the Python module for post-processing and analysis.

It is evident that Ithildin represents but one instance of software designed for the simulation of cardiac electrophysiology. Another FD solver is BeatBox [52], which also relies on domain splitting. A non-exhaustive list of examples of established simulation software based on the Finite Element Method (FEM) are Chaste [53], openCARP [54], lifex-ep [55], cbcbeat [56], GEMS [57], and CEPS [58]. Several of these packages have more advanced features than our software. In addition, some have broader applications than cardiac electrophysiology. More software and a benchmark study can be found in [59].

The main limitations of our software are the fact that Ithildin only supports the monodomain model and the fact that the domain decomposition happens solely in one coordinate direction. Additionally, it is maintained by a small research team. This paper serves to enhance the visibility of the software and to invite fellow cardiac modelers to use and contribute to the project. We believe that the GitLab environment is an effective medium for facilitating interaction between users, identifying issues and suggesting improvements.

## 7 Conclusion

In this work, we introduced Ithildin, an open-source library that allows for efficient numerical simulation and analysis of rotor waves. We demonstrated the versatility of Ithildin through a series of simulations, including spiral break-up in the Smooth-Karma model, the S1S2 protocol in the AP96 model, and 2D and 3D spiral waves in the BOCF model in ventricular geometry.

Our simulations highlighted several key features of Ithildin, such as the different implemented geometries and reaction terms, inhomogeneities, and stimuli, as well as recording data such as the pseudo-EGM or filament trajectories. These findings contribute to the growing understanding of rotor waves in cardiac electrophysiology and have the potential to inform future experimental and theoretical studies.

Overall, our work demonstrates the power of Ithildin as a tool for studying complex wave patterns in cardiac tissue. We hope that this library will be useful to researchers seeking to better understand the dynamics of rotor waves and their implications for cardiac function.

## Data availability

The source code of version 3.4.0 of the Ithildin software implemented for this paper is publicly available at https://gitlab.com/heartkor/ithildin and has been archived on Zenodo (DOI: 10.5281/zenodo.10876183). This archive also contains the data generated by the simulations used throughout this paper. Documentation of the code is publicly available at https://heartkor.gitlab.io/ithildin/.

## Appendix

This section contains C++ code to run the four simulations that are used throughout this paper. An overview of their parameters is given in tbl. 4.

### Sim. 1: Spiral break-up in the Smooth-Karma model

This is the source code to the Ithildin simulation used as the main example throughout this paper.

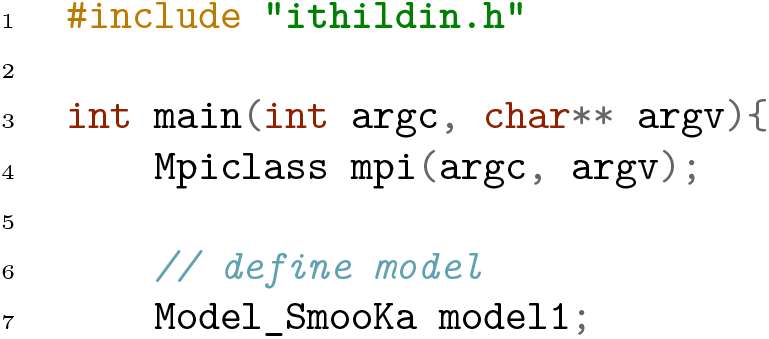
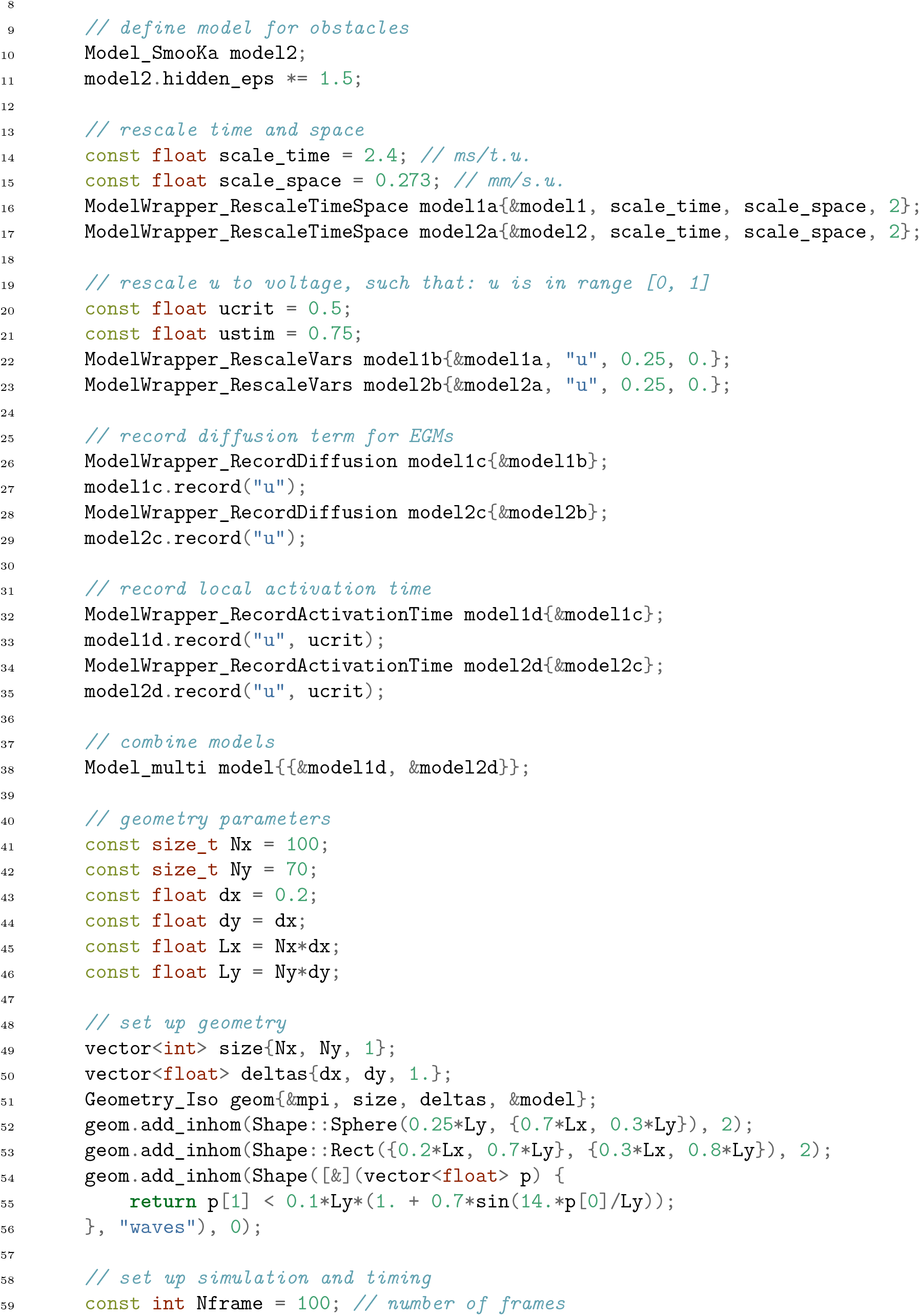
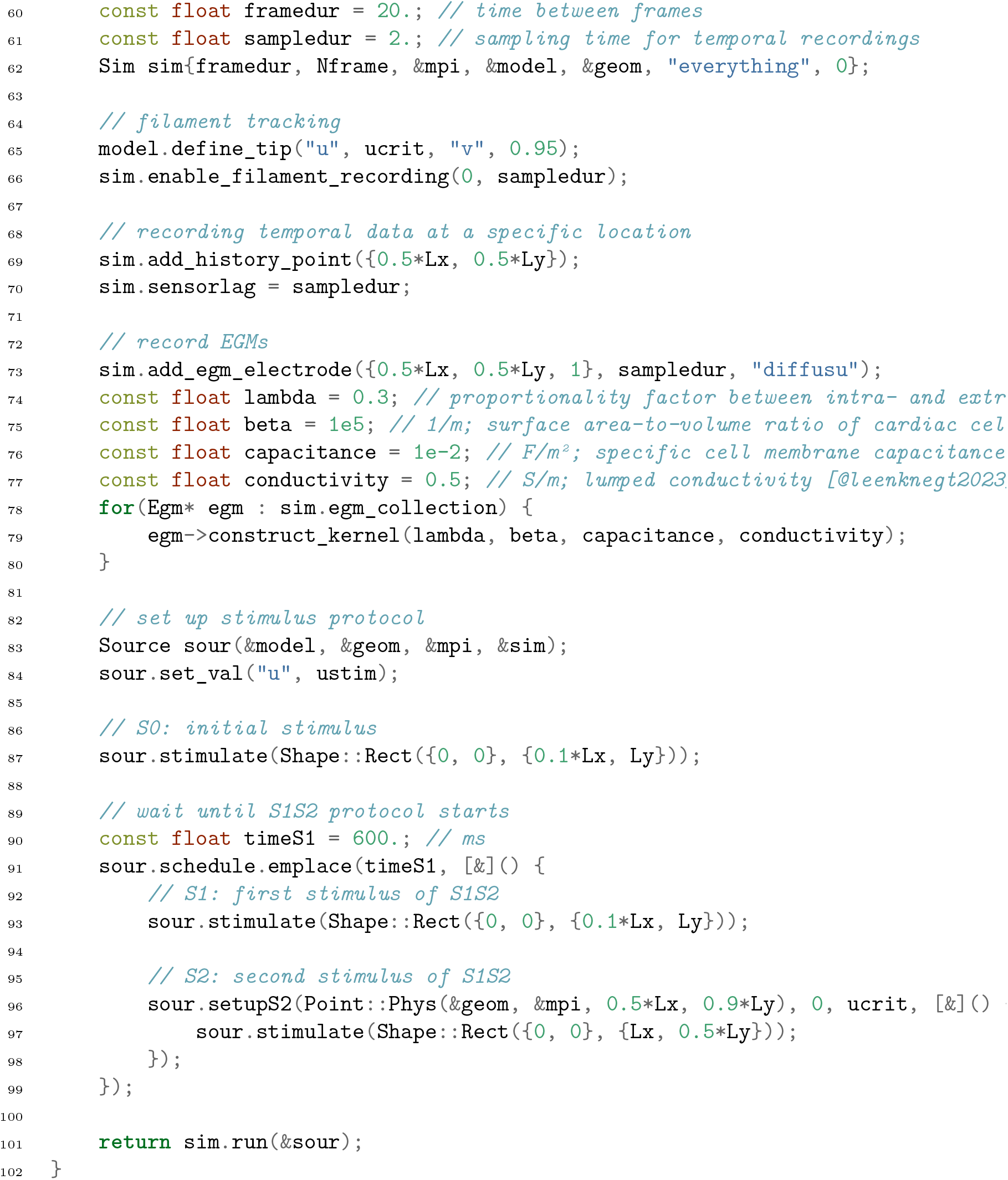

### Sim. 2: 2D spiral wave in the AP96 model

This example illustrates the S1S2 protocol in an easy-to-interpret setting.

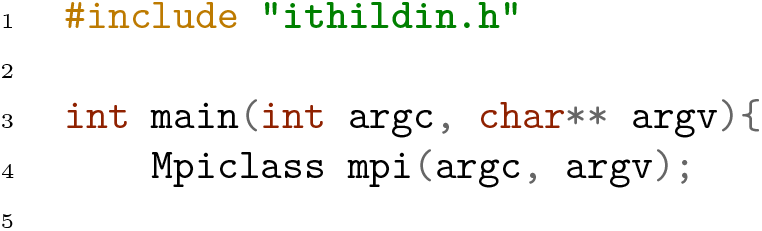
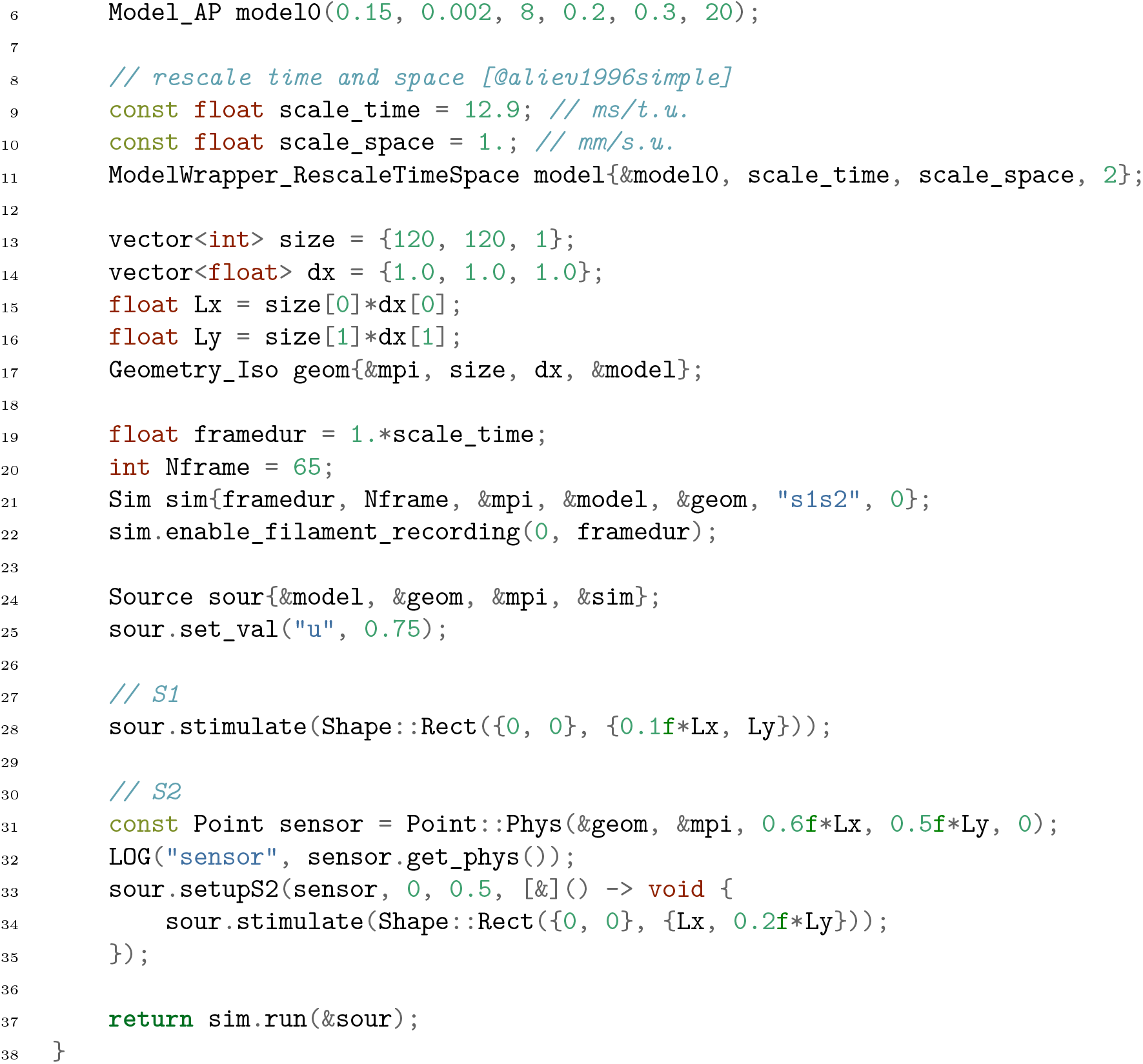

### Sim. 3: 2D spiral wave in the BOCF model

This simulation is used to get an initial state for a 3D simulation in ventricular geometry in the following example.

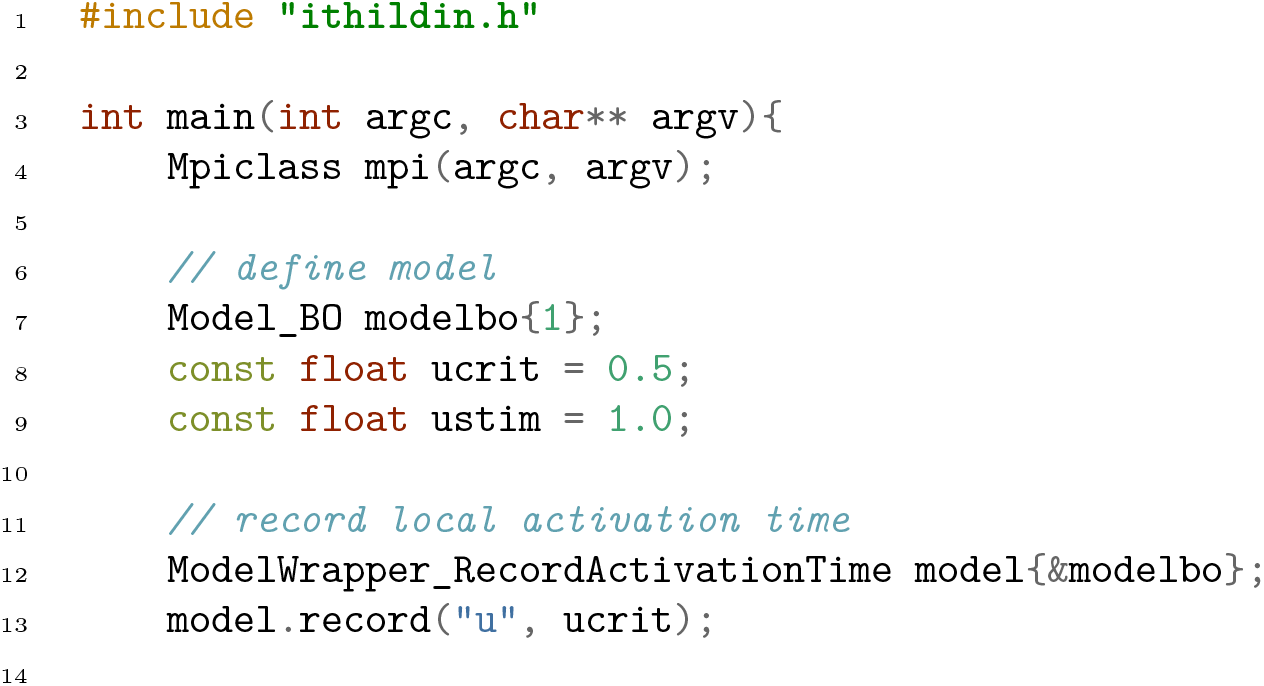
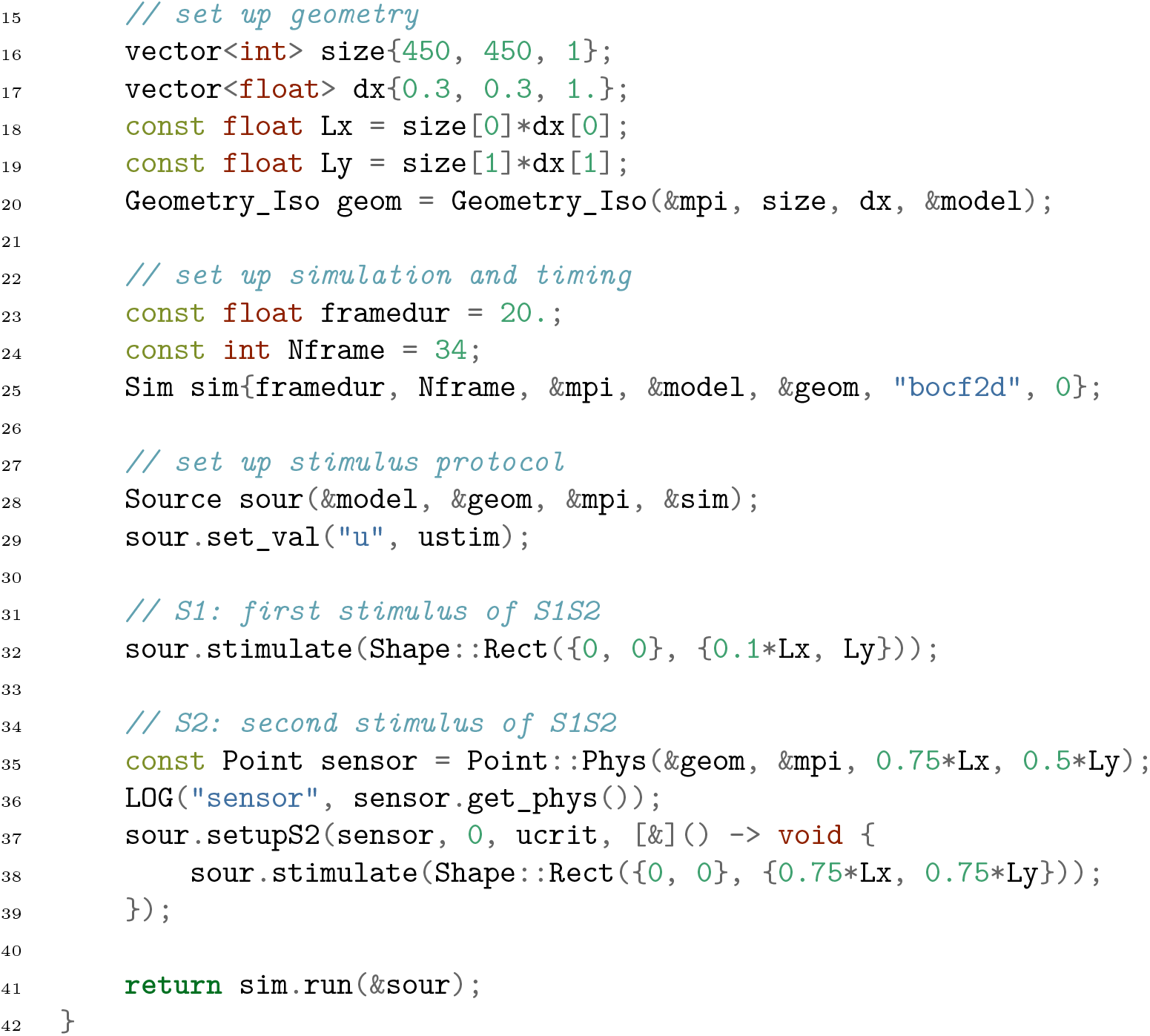

### Sim. 4: 3D spiral wave in the BOCF model

This 3D spiral wave is stimulated by extending a 2D spiral wave and placing it on ventricular geometry.

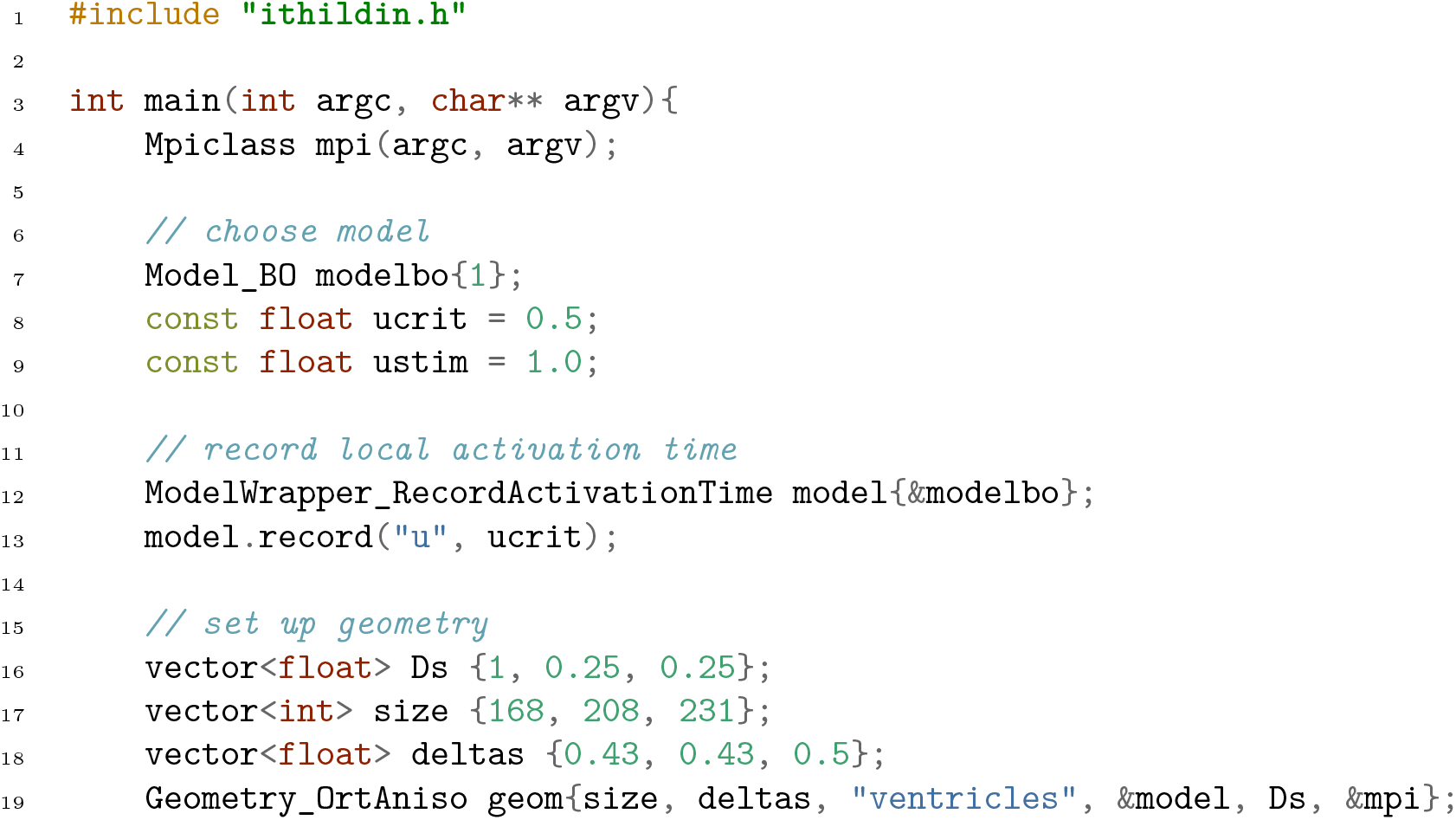
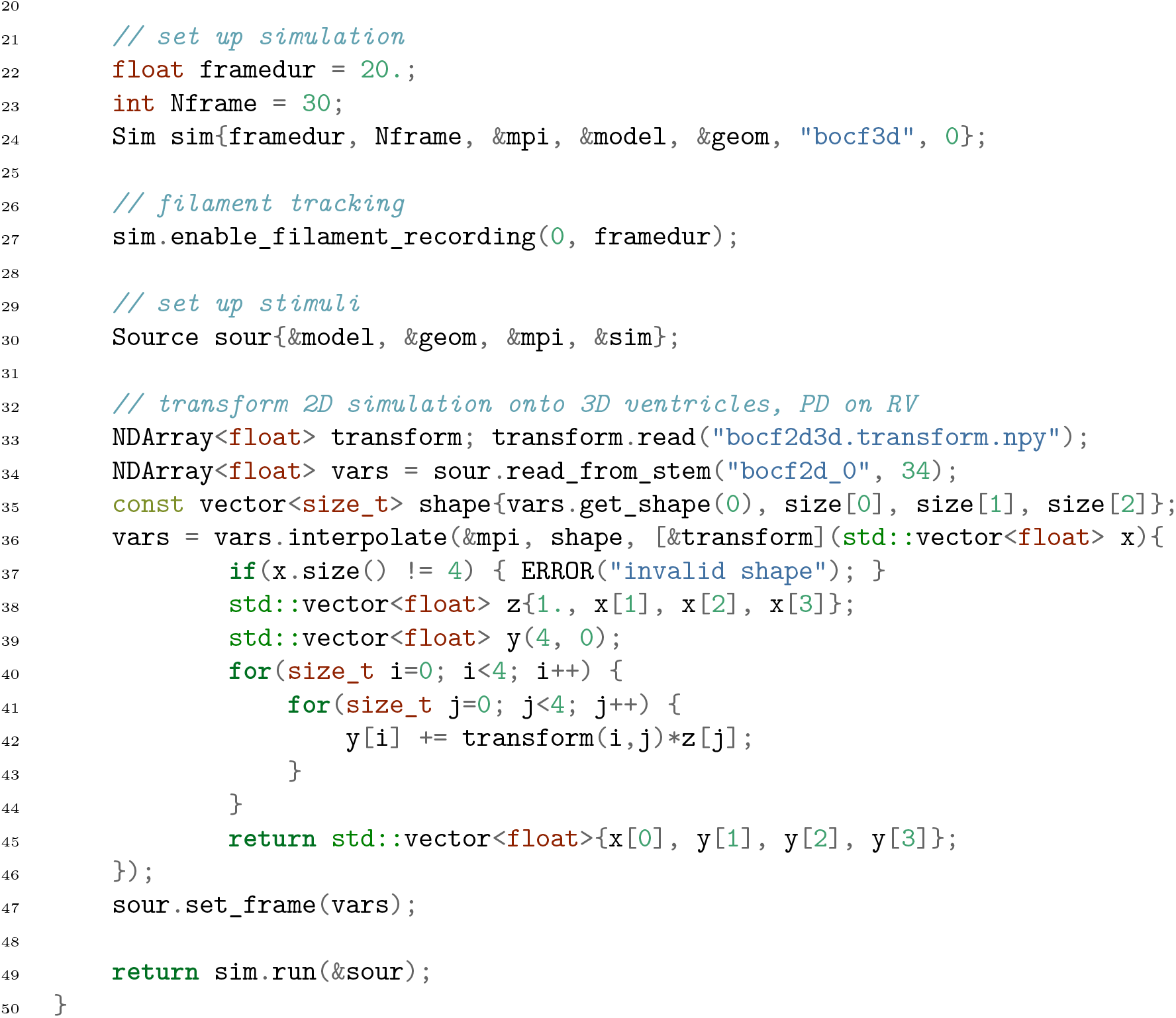

## Acknowledgments

We are grateful to Daniël A. Pijnappels, Antoine A.F. de Vries and our our other collaborators at the LUMC. Also, we would like to thank all members of Team HeartKOR.

While no generative AI has been used to write the Ithildin source code, generative AI has been used by the authors to aid in the writing of the text, specifically the Large Language Model (LLM) implementations GitHub Copilot, ChatGPT using GPT-3.5, as well as Mistral 0.2 via the Ollama software. The authors confirm that they have followed the current ethical publishing practices of PLOS One.

## Author contributions

**DK:** Conceptualization, Methodology, Software, Validation, Formal analysis, Investigation, Data Curation, Writing – original draft, Writing – review & editing, Visualization, Project administration. **MC:** Methodology, Software, Validation, Formal analysis, Investigation, Data Curation, Writing – original draft, Writing – review & editing, Visualization. **CZ:** Software. **OB:** Software. **HD:** Conceptualization, Software, Resources, Writing – review & editing, Supervision, Project administration, Funding acquisition.

